# Astrocyte glucocorticoid receptors mediate sex-specific changes in activity following stress

**DOI:** 10.1101/2024.09.17.613499

**Authors:** Lewis R. Depaauw-Holt, Sarah Hamane, Sarah Peyrard, Benjamin Rogers, Stephanie Fulton, Anthony Bosson, Ciaran Murphy-Royal

**Author notes:** **Corresponding author**: Ciaran Murphy-Royal CRCHUM, 900 rue, St. Denis, Montréal, Québec, Canada, H2X 0A9.

## Abstract

Interactions between orexin neurons and astrocytes in the lateral hypothalamus influence activity levels including circadian and motivated behaviour. These behaviors are disrupted by stress in rodents and form a hallmark of stress-related neuropsychiatric disorders. Here we set out to understand how stress influences activity and the underlying cellular mechanisms. We report that the long-term effects of stress on activity levels correlate with spontaneous firing of orexin neurons with hyperactivity in males and hypoactivity presented by female mice. These neuronal changes were accompanied by extensive astrocyte remodelling. Causal manipulations identified lateral hypothalamic astrocytes as key regulators of activity patterns. In the context of stress, genetic deletion of glucocorticoid receptors in lateral hypothalamic astrocytes rescued the effects of stress on orexin neuron firing, restoring activity to control levels in both males and females. Overall, these data suggest that astrocytic regulation of orexin neuron firing enables the maintenance of activity levels, and their dysfunction drives stress-induced activity dysregulation.

**Graphical Abstract:** 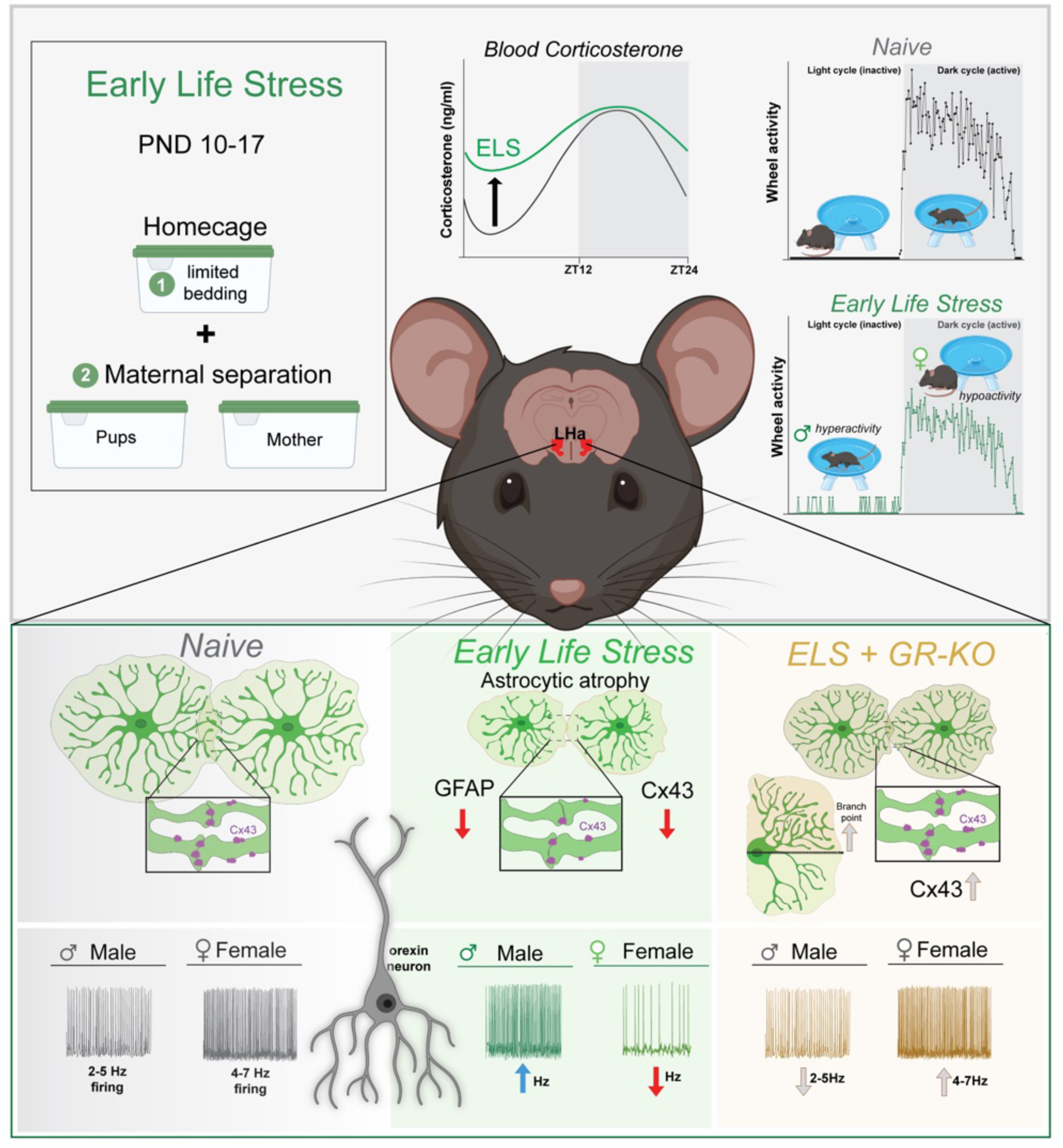

## Introduction

Astrocytes regulate multiple primary functions in the CNS^1^ and have coalesced with neurons to form an important element of the tripartite synapse^2^. Astrocyte morphology is complex and dynamic, permitting intimate contacts with diverse brain cells^3^. These varied contact points enable astrocytes to participate in the regulation of synaptic transmission and plasticity ^4,5^, central metabolism^6^, and homeostasis among other processes^7,8^. Across experimental models, astrocytes display a high level of heterogeneity throughout the developing and ageing brain^9,10^. During post-natal development astrocytes have been implicated in regulating the duration of “critical periods” by fine tuning excitability of underlying neuronal circuits^11^ and themselves display morphological and transcriptional phenotypes that coincide with important developmental milestones^12,13^. Morphological alterations in astrocyte processes stand as a defining corollary in pathological brain states^14–16^. One hallmark of the astrocytic response to insult, stressors, and disease includes changes in cell volume, evidenced by cellular atrophy or hypertrophy depending on the pathological state^17–23^. These volumetric aberrations are commonly associated with more discrete changes at the level of the peri-synaptic astrocyte process that is critical for regulating neurotransmitter^24–27^ and ionic homeostasis^28–33^ thereby modulating synaptic strength^34–37^. Not surprisingly, alterations in astrocytic morphology have been shown to influence behaviours including learning and memory^38^ and compulsive behaviors^3940^

In the context of stress, astrocyte dysfunction has been shown to drive impairments in synaptic transmission and plasticity^17^, with long-lasting effects on learning and memory^40^. Stress blunts synaptic plasticity by uncoupling the astrocytic metabolic network, creating an energetic substrate deficit in neighbouring neuronal compartments^17^. Stress, however, results in swath of behavioural phenotypes including changes in sleep-wake^41,42^, vigilance states^43,44^ and activity patterns across the day^45^, as well as motivated behaviors^46,47^. One key brain hub that has been implicated in these behavioral processes is the lateral hypothalamus (LH), containing orexin (hypocretin) neurons^48–51^. Orexin neuron firing is tightly controlled by multimodal mechanisms^52–54^ including local tuning by astrocytes^55,56^. There is a strong body of work supporting a role for the astrocyte neuron lactate shuttle (ANLS) in fueling orexin neuron firing with genetic perturbations of astrocyte function resulting in an instability of wakefulness^52,55,56^. This led us to question whether astrocyte dysfunction in the LH directly contributes to stress-induced behavioural changes across the light-dark cycle.

Here, using both female and male mice, we test the effect of an early-life stressor on lateral hypothalamic astrocytes and their control of region dependent behaviour. We found that early-life stress induced long-term changes in blood glucocorticoid levels and altered activity levels in a sex-specific manner with reduced activity in female and increased activity in male mice. These endocrine and behavioural changes were associated with extensive remodelling of astrocyte morphology, with distinct effects on orexin neuron firing patterns between sexes. By targeting glucocorticoid signalling specifically in LH astrocytes, we could abrogate the cellular, synaptic, and behavioural effects of early-life stress in both male and female mice. In conclusion, we reveal astrocytes as primary drivers of early-life pathological stress on neural activity and behaviour.

## Results

### Early Life Stress (ELS) elevates *nadir* blood glucocorticoids and perturbs diurnal activity rhythms

We leveraged a rodent model of ELS, including maternal separation (4hrs / day) between postnatal days (P)10-17 in combination with reduced bedding and nesting materials in all cages (**Fig. 1A,B**). Our previous work demonstrated this model of ELS induces a latent increase in blood glucocorticoids at P45^40^. We first aimed to replicate this finding and further examine whether the hormonal response was a consequence of a modified circadian set point. Consistent with previous data, we observed a significant rise in blood corticosterone in ELS mice at ZT2 (nadir) but not at ZT14 (zenith) (**Fig. 1C**). Like many other hormonal messengers, glucocorticoids are tightly regulated across the light-dark cycle^57,58^, and chronically high levels are associated with altered metabolism^59^, immune dysfunction^60^, and sleep-wake perturbation^61,62^. Interestingly, both chronic stress and chronic delivery of exogenous corticosterone to rodents disrupts sleep patterns^63^ and home cage running wheel activity^64,65^, a behaviour strongly regulated in a circadian and diurnal manner. To test, whether this ELS-induced hormone response was associated with changes in activity levels we single-housed mice from naïve and ELS litters into cages with *ad libitum* access to a running wheel for two weeks (1 week habituation, 1 week data analysis). We observed a significant reduction in running wheel activity in ELS mice as measured by distance ran per day (**Fig. 1D-F**). This effect was due to an altered activity rhythm (**Fig. 1F**), where ELS mice exhibited hyperactivity during the light cycle (ZT0-12) and hypoactivity during the dark cycle (ZT12-24) resulting in diminished AUCs independent of peak running wheel activity (**Fig. 1G-J**). We next partitioned the data by sex and found a striking sex difference in the effect of ELS on running wheel activity (**Fig. 1K-N**). Male mice subjected to ELS displayed no change in the distance ran per day but displayed significant light-cycle hyperactivity compared to naïve males (**Fig. 1K&M**). The inverse was true for female ELS mice, with a significant reduction in the distance ran per day with no change in light-cycle activity, exhibiting a behavioural phenotype isolated to the wake period (**Fig. 1L&N, Supplemental Fig. 1D**). Importantly, we observed no significant effects of this ELS paradigm on total body weight at P45 (**Supplemental Fig.1A**) or any significant changes to body weight after the 2-week running wheel protocol (**Supplemental Fig.1B**) ruling out potential adverse effects on basal metabolism or metabolic adaptation to exercise. Additionally, this ELS paradigm has been shown to influence transcriptional profiles in reward centres of the brain^66–68^ suggesting our running wheel phenotype could be biased by a weakened hedonic drive in ELS mice. To answer this question, we subjected a cohort of ELS and naïve mice to a sucrose preference test (SPT) but saw no influence of stress on sucrose preference in either sex, ruling out anhedonia as a confounding factor in our experiment (**Supplemental Fig.1C**). These data reveal that ELS elevates stress hormones in a diurnal manner and has a sexually dimorphic effect on activity rhythms.

**Figure 1.**
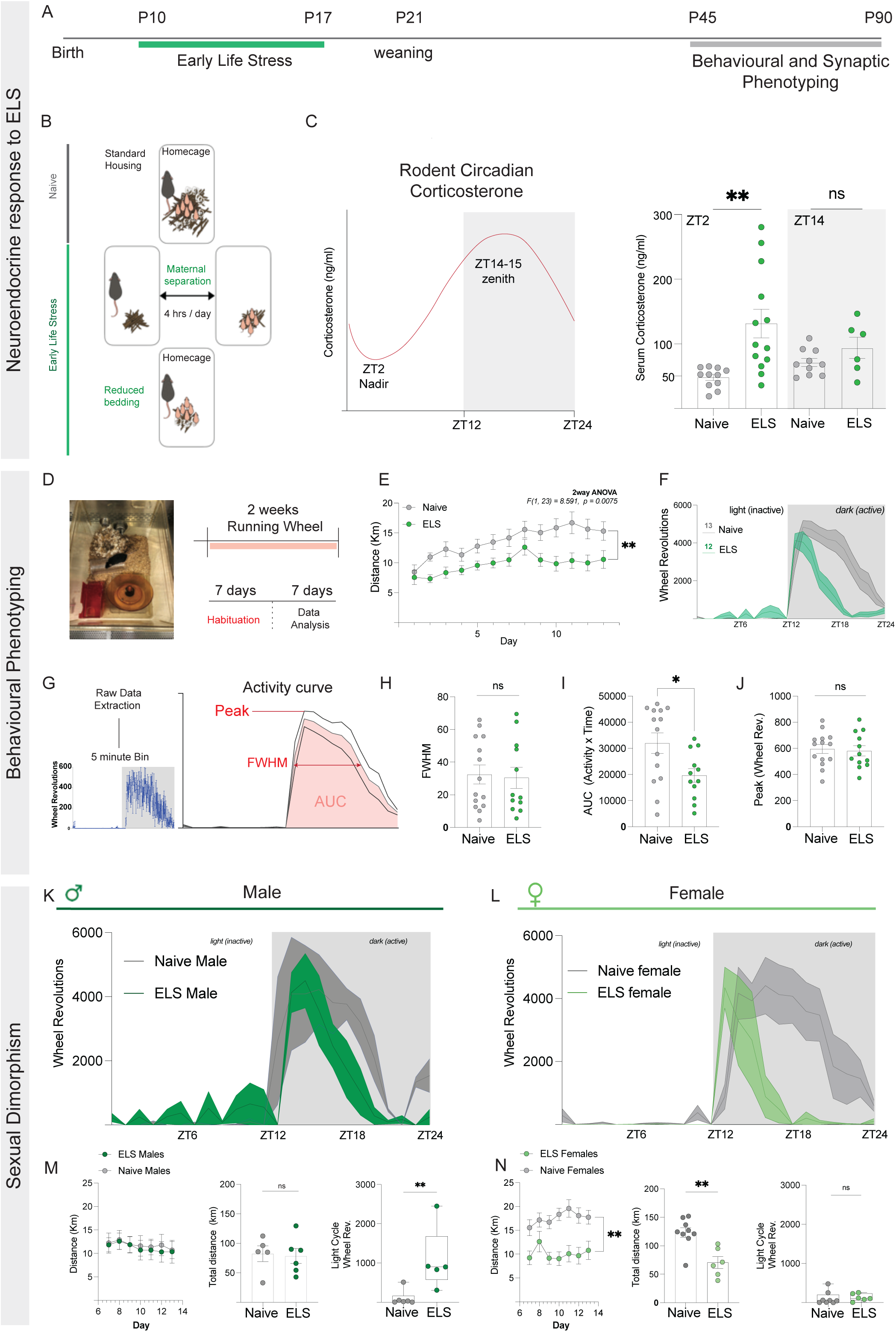
ELS induces elevations in blood corticosterone and differentially alters diurnal activity rhythms in male and female mice. **A**) Experimental timeline with ELS occurring between P10-P17. **B**) ELS paradigm; maternal separation 4 hours/day and 70% reduced bedding in home cage and separation cages. **C**) Circadian rhythm of rodent corticosterone (nadir = ZT2, zenith = ZT14-15). Serum corticosterone measurements significantly elevated at ZT2 (unpaired *t*-test: naïve = 48.04 ng/ml (N=11), ELS = 131.6 ng/ml (N=13), *p*=0.0026) but not ZT14 (unpaired *t*-test: naïve = 71.21ng/ml (N=10), ELS =93.75 ng/ml (N=6), *p*=0.154). **D**) Running wheel protocol; 2-week single housing with *ad libitum* access to horizonal running wheels. **E**) Distance ran per day of naïve (N=13) and ELS (N=12) mice (2way ANOVA; *F*(1,23) = 8.591, *p*=0.0075). **F**) representative 24-hour trace of running wheel activity from naïve and ELS mice. **G**) Analyses of 24-hour running wheel activity curves from naïve and ELS mice. **H**) FWHM (Mann-Whitney test: naïve = 32.44, ELS = 30.41, *p*=0.849). **I**) AUC (Mann-Whitney test: naïve = 31987, ELS = 19649, *p*=0.031). **J**) Peak wheel activity (unpaired *t*-test: naïve =595.8, ELS =580.6, *p*=0.773). **K**) 24-hour traces from naïve and ELS male mice. **L**) 24-hour traces from naïve and ELS female mice. **M**) Distance ran per day (2way ANOVA; *F*(1,9) =0.046, *p*=0.834), Total distance ran (Mann-Whitney test: naïve =82.40, ELS =78.42, *p*=0.9307), and light-cycle activity counts (Mann-Whitney test: naïve =23.83, ELS = 898.7, *p*=0.0087) for naïve and ELS male mice. **N**) Distance ran per day (2way ANOVA; *F(*1,13) =15.63, *p*=0.0016), Total distance ran (Unpaired *t*-test: naïve =123.3, ELS =70.64, *p*=0.0016), and light-cycle activity counts (Mann-Whitney test: naïve =50.17, ELS =108.8, *p*=0.282) for naïve and ELS female mice. N= mice. Ns, not significant, *P < 0.05, **P < 0.01.

### ELS alters morphology of lateral hypothalamic astrocytes

Running wheel activity is entrained by multiple nuclei from a variety of brain regions^69–71^ including the lateral hypothalamus (LH)^72,73^. The LH is a hub that integrates circadian inputs^74^, metabolic cues^75–77^, and local synaptic inputs^78,79^ to drive activity. Orexin neurons of the LH are wake promoting, and as such display higher activity during the onset of the dark cycle and are less active during the light cycle to allow for sleep onset and maintenance respectively^56^. Prior work has shown that astrocytic metabolic networks sustain orexin neuron activity to drive sleep-wake cycles^55,56^. Considering the potential impact of early-life stress on neural circuit development, we first conducted immunohistochemistry (IHC) on coronal brain slices of the LH from naïve and ELS mice and quantified the number of Orexin-A positive neurons. We found no impact of ELS on the number of Orexin-A+ cells with no sex differences (**Fig. 2A**). We next quantified the number of astrocytes (s100β+) in the LH and, again, observed no influence of stress (**Fig. 2B**), suggesting our ELS paradigm does not hinder post-natal differentiation of orexin neurons and astrocytic neighbours.

**Figure 2.**
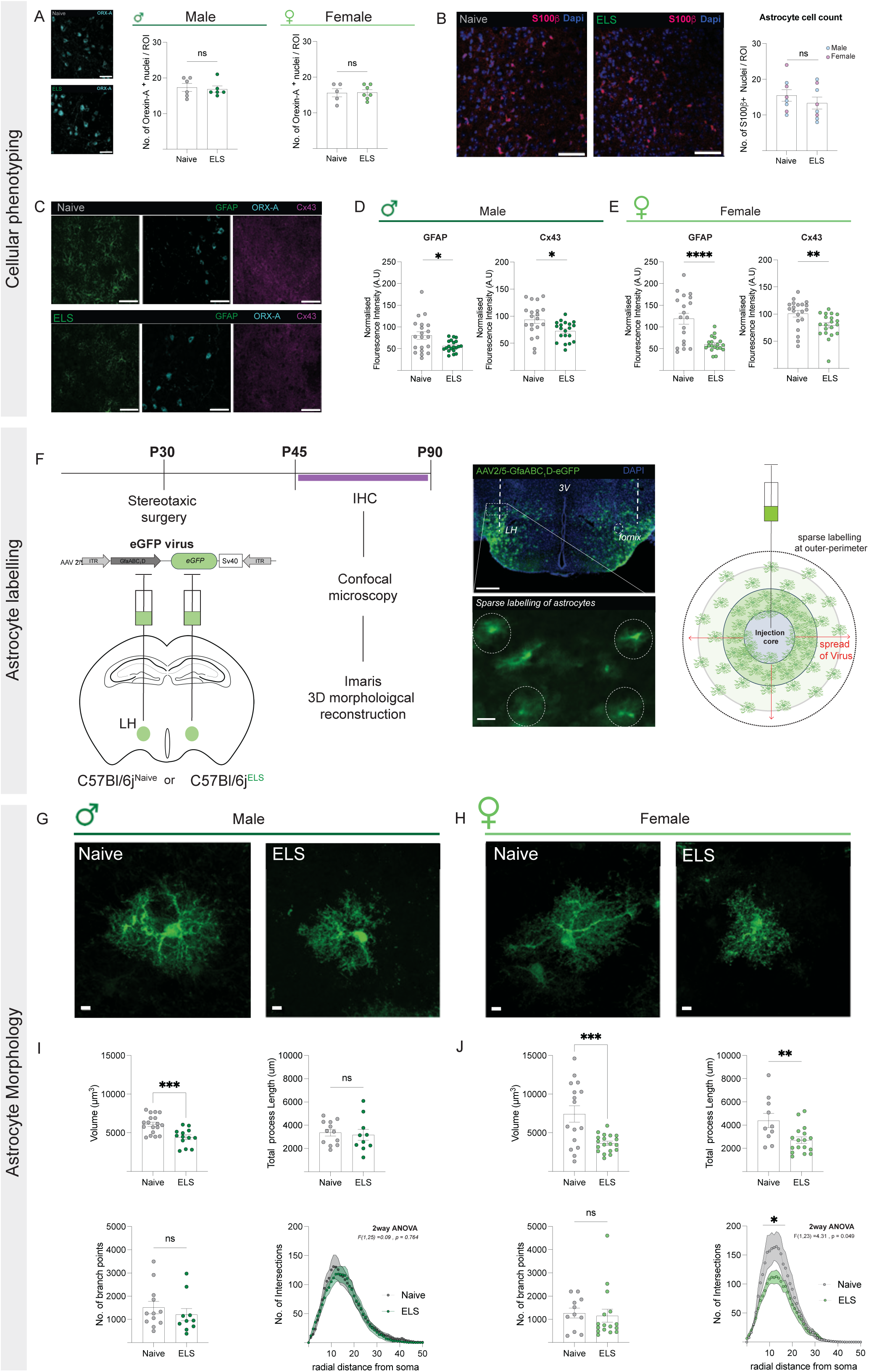
ELS modifies astrocyte morphology in the LH. **A**) Immunohistochemistry (IHC) of Orexin-A positive cells in the LH. Quantification of Orexin-A+ cells between naïve and ELS male mice (unpaired *t-*test: naïve = 17.33 (N=6), ELS = 16.83 (N=6), *p*=0.727) and naïve and ELS female mice (unpaired *t*-test: naïve = 15.60 (N=5), ELS = 15.71 (N=7), *p*=0.933). **B**) Quantification of s100β+ astrocytes in the lateral hypothalamus (unpaired *t*-test: naïve = 14.50 (N=8), ELS =12.00 (N=8), *p=*0.367). **C**) Representative IHC of GFAP, Orexin-A, and Cx43, In the LH of naïve and ELS mice. **D**) Normalised fluorescence intensity (a.u) measures of GFAP and Cx43 in the LH of naïve and ELS male mice (GFAP - unpaired *t*-test: naïve = 80.33 (N=4, n=20 cells), ELS = 55.91 (N=4, n=20 cells), *p*= 0.011. Cx43 - unpaired *t*-test: naïve = 93.74 (N=4, n=20 cells), ELS =72.68 (N=4, n=20 cells), *p*=0.011). **E)** Normalised fluorescence intensity (a.u) measures of GFAP and Cx43 in the LH of naïve and ELS female mice (GFAP - unpaired *t*-test: naïve =119.2 (N=4, n=19 cells), ELS =58.87 (N=4, n=19 cells), *p*<0.0001, Cx43 - unpaired *t*-test: naïve = 100.8 (N=4, n=20 cells), ELS =79.27 (N=4, n=20 cells), *p*=0.009). **F**) Experimental timeline and viral constructs for sparse labelling of astrocytes in the LH. Scale bars = 500um (top) and 25um (bottom). **G-H**) Representative AAV-GfaABC_1_D-eGFP expressing astrocytes in the LH from naïve and ELS mice, scale bars = 5um. **I**) Morphological reconstruction of LH astrocytes from naïve and ELS male mice: Volume (unpaired *t*-test: naïve =5916 (N=4, n=18 cells), ELS =4430 (N=3, n=13 cells), *p*=0.0004), Total process length (unpaired *t*-test: naïve =3365 (N=3, n=12 cells), ELS =3168 (N=3, n=10 cells), *p*=0.717), No. of branch points (Mann-Whitney test: naïve=1256 (N=3, n=12 cells), ELS =943 (N=3, n=10 cells) *p*=0.314), Sholl analysis (2way ANOVA; F(1,25) =0.09, *p*=0.764). **J**) Morphological reconstruction of LH astrocytes from naïve and ELS female mice: Volume (unpaired *t*-test: naïve =7740 (N=4, n=16 cells), ELS =3659 (N=4, n=19 cells), *p*=0.0006), Total process length (unpaired *t*-test: naïve =4396 (N=3, n=10 cells), ELS =2701 (N=4, n=17 cells), *p*=0.009), No. of branch points (Mann-Whitney test: naïve=1209 (N=3, n=11 cells), ELS =720.5 (N=4, n=16 cells) *p*=0.488), Sholl analysis (2way ANOVA; F(1,23=4.31, *p*=0.049). N= mice. Ns, not significant, *, p < 0.05; **, p < 0.01 and ***, p < 0.001.

Given the abundance of literature highlighting the acute and chronic effects of stress on astrocytes^17,80–84^ we then assessed the expression levels of two canonical markers for astrocytes, glial fibrillary acidic protein (GFAP); an intermediate filament protein, and connexin-43 (Cx43); one of the functional gap-junction channel proteins in astrocytes that is critical for metabolic support of orexin neuron firing^55^. (**Fig. 2C**). We observed significant reductions in GFAP and Cx43 in both male and female ELS mice (**Fig. 2D&E**). Changes in GFAP expression have long been implicated in astrocytic “reactivity states”, but their expression levels are restricted to large primary branching processes, which represents a minute portion of the astrocytic cytosol^19^. To overcome this, we injected mice with a low-titre AAV2/5-GfaABC_1_D-eGFP virus to sparsely label astrocytes within the LH and conducted in-depth morphological analyses (**Fig. 2F-H**). Consistent with our IHC experiments, we observed a significant reduction in total three-dimensional volume of LH astrocytes in both male and female ELS mice (**Fig. 2I&J**). Interestingly, we note effects of ELS on total process length and ramification in female mice, but not in male mice (**Fig. 2I&J**). Taken together with our IHC data we report a significant reorganisation of astrocyte morphology in the LH of both male and female ELS mice in the absence of change to the number of neuronal and astrocytic cell types.

### Driving astrocyte Gq-coupled calcium signaling in the LH impairs activity rhythms

Experimental manipulation of orexin neuron activity has been shown to directly influence wakefulness^85,86^ and wheel running behaviours^72^. While ELS exerts profound effects on both diurnal activity rhythms and astrocytic morphology in the LH, it’s unclear whether astrocytes themselves participate in regulating running wheel activity across the light-dark cycle. To address this, we used a chemogenetic approach to specifically drive calcium activity in a temporally precise and brain-region specific manner. Compartmentalised fluctuations in astrocyte calcium have been shown to be a primary signaling mechanism by which astrocytes fine-tune neuronal output^39,87–91^. A cohort of naïve mice were injected with hM3D(Gq)- or hM4D(Gi)-coupled Designer Receptors Exclusively Activated by Designer Drugs (DREADDs) under the control of GfaABC_1_D promoter to selectively induce calcium fluctuations in LH astrocytes. To chemogenetically activate astrocytes, Clozapine-N-Oxide (CNO) was added to the drinking water for 3 days, standardised to water intake and body weight (**Fig. 3A**). Oral delivery of CNO was chosen to mitigate handling stress associated with intraperitoneal injection. Remarkably, we report a significant effect of CNO exclusively on hM3D(Gq)-injected mice and not hM4D(Gi)-injected or mCherry-injected controls (**Fig. 3B-D**). hM3D(Gq)-injected mice exhibited significant elevations in light-cycle wheel activity after administration of CNO (**Fig. 3B**). This isolated effect of hM3D(Gq)- and not hM4D(Gi)-activation was consistent at 3 ascending doses of CNO **(Supplemental Fig. 2**). These findings are consistent with recent data^39^ showing no effect of hM4D(Gi)-activation on behaviour in naïve mice^39^. Taken together, our data suggest that Gq-GPCR mediated fluctuations in astrocyte calcium and downstream signalling in the LH are important for regulating diurnal entrainment of running wheel activity.

**Figure 3.**
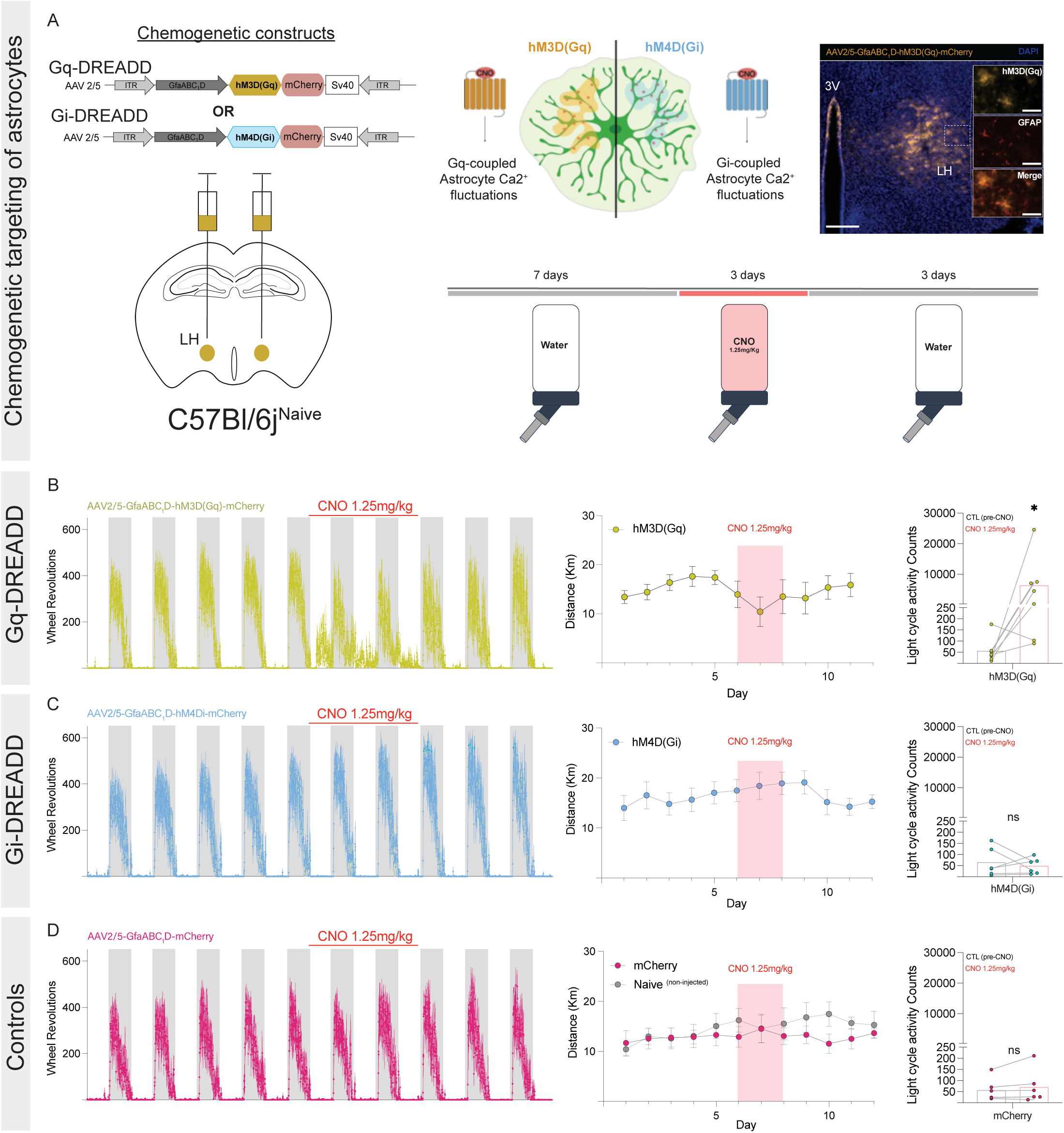
Gq-coupled calcium fluctuations in LH astrocytes perturbs diurnal activity rhythms in naïve mice. **A**) Chemogenetic constructs and expression in LH astrocytes, scale bars = 200um, and 50um. Chronic delivery of clozapine-N-oxide (CNO) in drinking water (1.25mg/kg). **B**) Representative traces of running wheel activity from hM3D(Gq)-injected mice (N=7), distance ran per day, and quantification of light cycle activity pre and during delivery of CNO (ratio paired *t*-test: *t*6=3.34, *p*=0.016). **C**) Representative traces of running wheel activity from hM4D(Gi)-injected mice (N=6), distance ran per day, and quantification of light cycle activity pre and during delivery of CNO (paired *t*-test: *t*5=0.574, *p*=0.591). **D**) Representative traces of running wheel activity from mCherry-injected mice (N=6), distance ran per day, and quantification of light cycle activity pre and during delivery of CNO (paired *t*-test: *t*5=1.33, *p*=0.241). N= mice. *P < 0.05.

### ELS modifies intrinsic firing properties of orexin neurons in a sexually dimorphic manner

We observed a clear sex difference in the behavioural response to ELS (**Fig. 1G&H**), with male light-cycle hyperactivity and female dark-cycle hypoactivity. Based on our cellular characterisation of ELS in the LH, this effect was not a consequence of altered neuronal population sizes, but rather a reorganisation of local astrocytic morphology (**Fig. 2A**). Given the longstanding importance of astrocytes in regulating neuronal excitability^92–97^ we next assessed whether these changes were reflected at the level of orexin neuron activity. To identify orexin neurons, we injected an AAV2/8-miniHCRT-tdTomato virus into the LH of naïve and ELS mice. After a minimum of 2 weeks, we prepared acute brain slices and performed whole-cell patch clamp recordings of tdTomato+ cells in the peri-fornical area of the LH (**Fig. 4A&C**). We first validated the specificity of the AAV2/8-miniHCRT-tdTomato construct and found an average of 91.07% (± 4.49) of tdTomato+ cells colocalized with Orexin-A (**Fig. 4B**). Orexin neurons have unique electrophysiological signatures including low-threshold spikes after depolarising currents, inward rectification in response to hyperpolarising currents (I_h_-current), and intrinsic firing rates of 2-4 Hz in the presence of 2.5mM of glucose^55,98,99^. We first compared our tdTomato+ cells to biocytin filled orexin neurons, identified post-hoc through IHC (**Supplemental Fig.3A**). We report no significant differences in firing rates or resting membrane potentials (RMP) of these neurons, demonstrating no adverse effects of AAV2/8-miniHCRT-tdTomato expression on intrinsic membrane properties.

**Figure 4.**
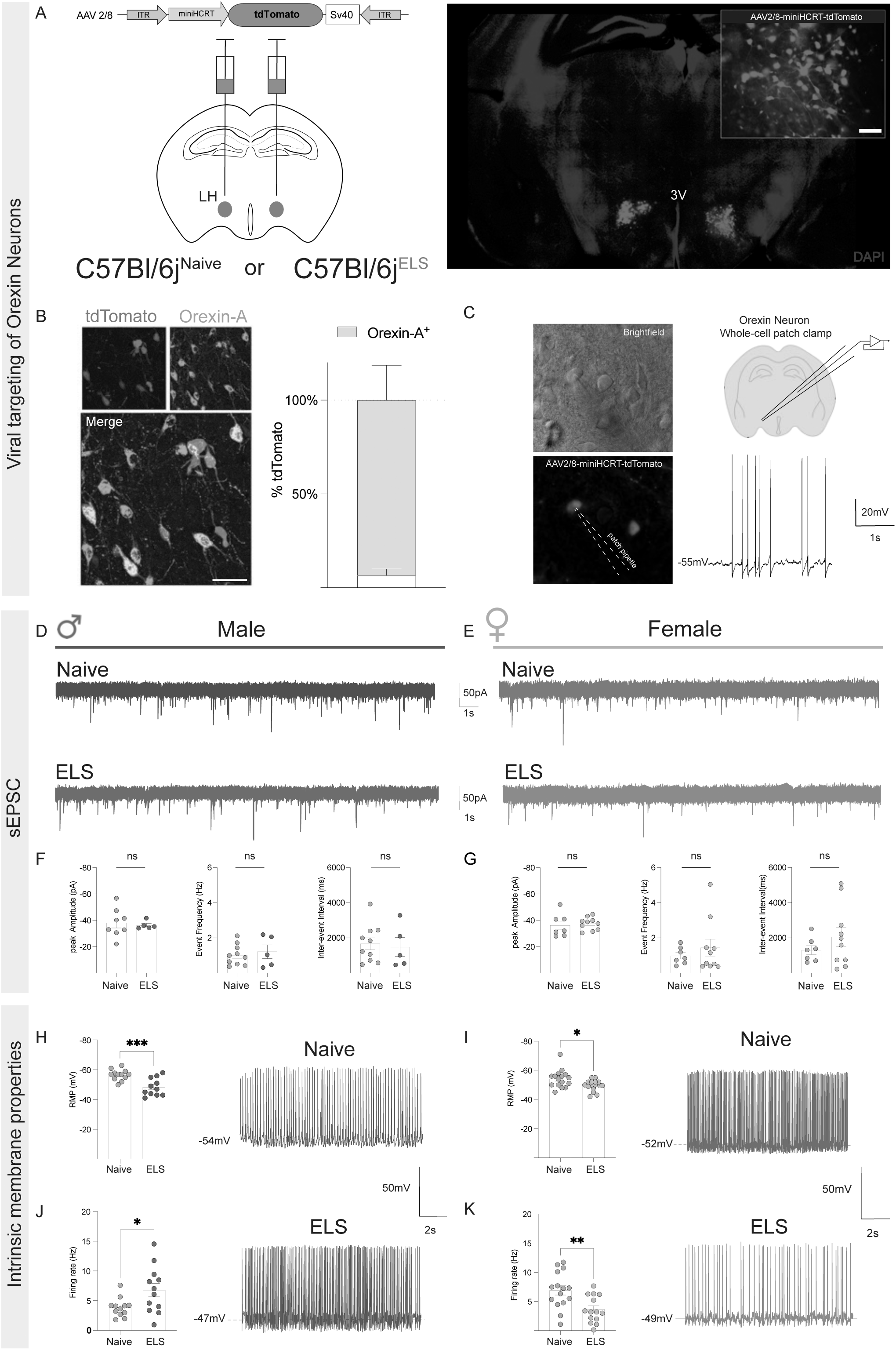
ELS modifies intrinsic firing rates of orexin neurons in a sex specific manner. **A**) Viral targeting of orexin neurons in the LH, scale bar = 25um **B**) Validation of AAV-2/8-miniHCRT-tdTomato construct, scale bar =50um. **C**) Representative neurons from acute slice preparation under brightfield and mCherry filter cube fluorescence with representative spontaneous firing rates of 3-4Hz. **D-E)** Representative traces of spontaneous EPSCs (sEPSC) from orexin neurons in voltage-clamp (-70mV) configuration from naïve and ELS mice. **F**) Male mice orexin neuron sEPSC peak amplitude (unpaired *t*-test: naïve =-38.14 (N=3, n=8 cells), ELS =-36.35 (N=2, n=5 cells), *p*= 0.728), event frequency (unpaired *t*-test: naïve =1.022 (N=3, n=10 cells), ELS =1.215 (N=2, n= 5 cells), *p*=0.616), inter-event interval (unpaired *t*-test: naïve=1665 (N=3, n=10 cells), ELS =1478 (N=2, n=5 cells), *p*=0.765). **G**) Female mice orexin neuron sEPSC peak amplitude (unpaired *t*-test: naïve = -37.57 (N=3, n=7 cells), ELS = -36.31 (N=3, n=10 cells), *p*=0.705), event frequency (Mann-Whitney test: naïve = 0.836 (N=3, n=7 cells), ELS = 0.853 (N=3, n= 10 cells), *p*=0.0.720), inter-event interval (unpaired *t*-test: naïve = 1306 (N=3, n=7 cells), ELS = 2049 (N=3, n=10 cells), *p*=0.303). **H**) Resting membrane potentials (RMP) of orexin neurons from naïve and ELS male mice (unpaired *t*-test: naïve =-56.65 (N=3, n= 12 cells), ELS = -48.36 (N=4, n=11 cells), *p*=0.0005). **I**) Resting membrane potentials (RMP) of orexin neurons from naïve and ELS female mice (unpaired *t*-test: naïve =-54.19 (N=4, n= 16 cells), ELS = -49.66 (N=4, n=16 cells), *p*=0.0169). **J**) Spontaneous firing rates of orexin neurons from naïve and ELS male mice (unpaired *t*-test: naïve = 3.85 (N=4, n= 12 cells), ELS =6.789 (N=4, n=12 cells), *p*=0.025) **K**) Spontaneous firing rates of orexin neurons from naïve and ELS female mice (unpaired *t*-test: naïve = 6.91 (N=4, n= 15 cells), ELS = 3.648 (N=4, n=14 cells), *p*=0.0037). N= mice . Ns, not significant, *, p < 0.05; **, p < 0.01 and ***, p < 0.001.

We next conducted voltage-clamp recordings of orexin neurons from naïve or ELS brain slices to examine the effect of ELS on spontaneous excitatory post-synaptic currents (sEPSC) (**Fig. 4D&E**). To our surprise we observed no significant differences in sEPSC measures; frequency, peak amplitude, or inter-event interval (**Fig. 4F&G**). We then assessed intrinsic membrane properties of orexin neurons and found a significant depolarisation of the RMP of orexin neurons from both male and female ELS mice (**Fig. 4H&I**). Furthermore, when we measured intrinsic firing rates we observed a striking sex difference in response to ELS. Male ELS mice displayed elevated firing rates compared to naïve males (**Fig. 4J**), whereas firing rates were greatly attenuated in female ELS mice (**Fig. 4K**). Of note, we also report a sex difference in naive conditions, where female orexin neurons exhibit higher firing rates (5-7 Hz) compared to male litters (2-4Hz) (**Supplemental Fig.3B**). While there is a small body of literature demonstrating sex differences in orexin excitability with respect to mRNA expression (increased expression of c-fos)^100^, this, to our knowledge, is the first demonstration of sex differences in basal orexin neurons firing. Given the wake-promoting nature of orexin neurons, these results are consistent with our behavioural phenotypes, where male ELS mice displayed hyperactivity and elevated firing rates, and female ELS with hypo-activity and attenuated firing rates. Additionally, we observed no change in other measures of intrinsic excitability such as input resistance (R_in_) in neither male nor female mice (**Supplemental Fig. 3C&D**). These data demonstrate the sexual dimorphic effects of ELS on behaviour is reflected at the synaptic level in the LH.

### Astrocyte glucocorticoid receptor KO rescues the cellular & behavioral effects of ELS

So far, we have shown that ELS impacts endocrine function (**Fig. 1C**), astrocyte morphology (**Fig. 2F-H**), and orexin neuron firing (**Fig. 4 J-K**) culminating in behavioural dysfunction (**Fig. 1K-N**). The mechanistic link between these diverse cellular, synaptic, and behavioural observations however, has yet to be addressed. Prior work demonstrated astrocytes are highly sensitive to stress-induced fluctuations in glucocorticoids with glucocorticoid signalling suggested to directly impact gap-junction coupling in these cells^17^. We conducted further IHC staining against glucocorticoid receptors (GR) in the LH and found a significant increase in nuclear localisation of GRs in astrocytes (**Supplemental Fig. 4A&B**), a proxy of GR signalling as nuclear translocation of GR occurs with their activation. Given we observed a modest (∼20%) but significant reduction in connexin-43 fluorescence in ELS astrocytes (**Fig. 2D&E**) accompanied by elevated blood corticosterone in ELS mice (**Fig.1C**), we questioned whether local suppression of glucocorticoid signalling in LH astrocytes could abrogate the effects of ELS on cellular, synaptic and behavioural scales.

To determine the contribution of astrocytic GR signalling in mediating the effects of stress we targeted this pathway in a cell-type specific and brain region specific manner. The selective deletion astrocytic GRs in the LH was achieved using a transgenic mouse line with a floxed Nuclear Receptor Subfamily 3 Group C Member 1 (*Nr3c1*^fl/fl^) allele. *Nr3c1*^fl\fl^ mice were bilaterally injected with AAV2/5-GfaABC_1_D-Cre-eGFP to delete GRs in astrocytes exclusively in the LH [GR^KO^] (**Fig. 5A**). Stereotaxic surgery was completed at P30 (13 days after ELS) and Cre-injected mice were compared to both ELS *Nr3c1*^fl\fl^ and naïve *Nr3c1*^fl\fl^ mice injected with a control virus (AAV2/5-GfaABC_1_D-eGFP). Viral injections did not influence neuronal (Orexin-A+) or astrocyte (s100β+) cell counts (**Fig. 5B**), and cre-recombination was restricted to astrocytes with an average specificity of 94.5% (± 4.41) in s100β+ astrocytes and expression efficiency of 88.2% (± 5.61) (**Fig. 5B, Supplemental Fig. 5D**). Compared to control astrocytes (Cre-negative) within the same brain and eGFP-control astrocytes from injected littermates, we observed a significant reduction in nuclear GR fluorescence (**Supplemental Fig. 5A-C**).

**Figure 5.**
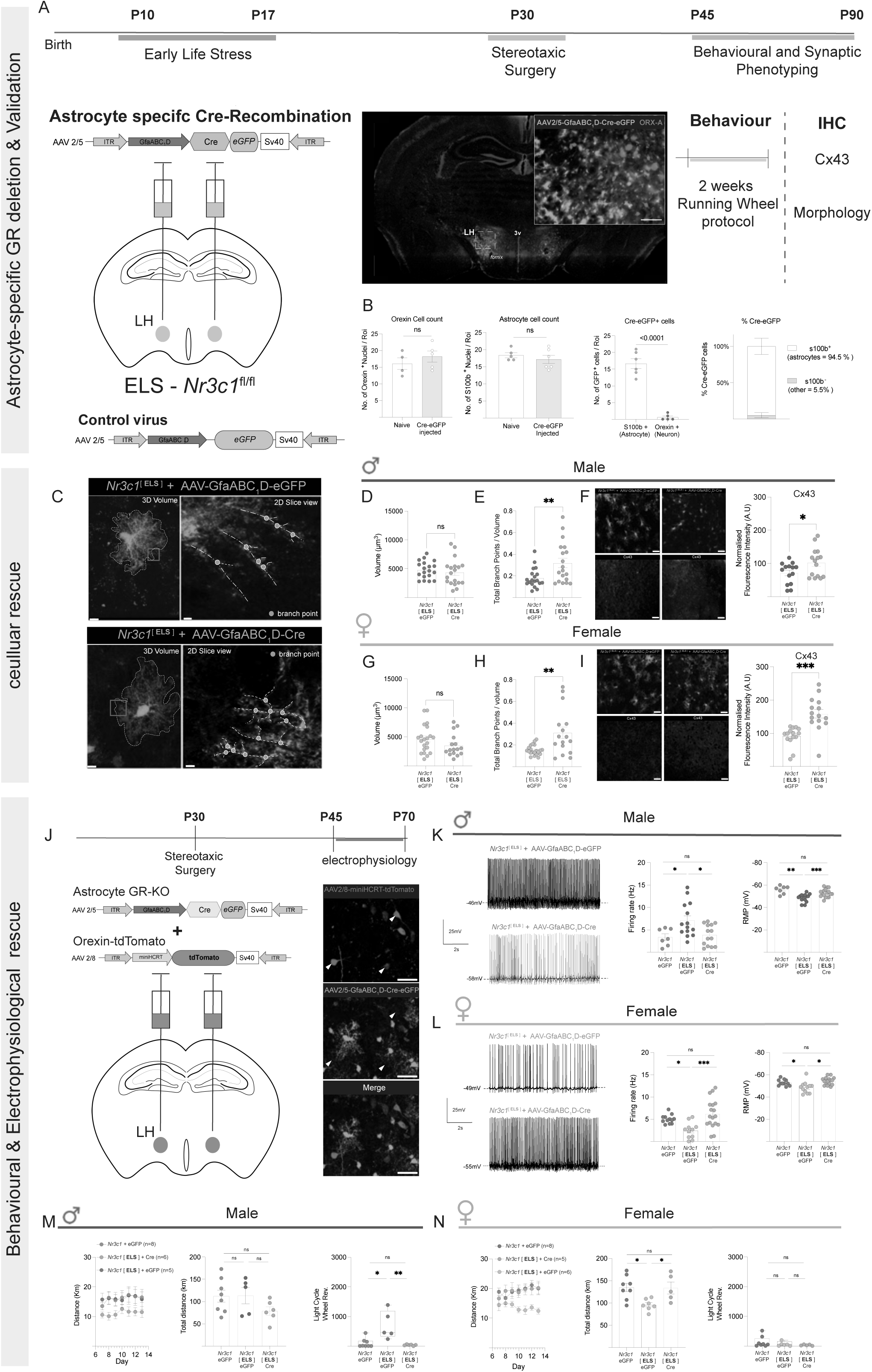
Astrocyte specific deletion of glucocorticoid receptors (GR) in the LH corrects cellular and behavioural impairments induced by ELS. **A**) Experimental timeline for combined ELS and genetic deletion of astrocyte GRs in the LH. Separate cohorts of mice subjected to behavioural protocols or IHC, scale bar = 50um. **B**) Validation of AAV2/5-GfaABC_1_D-Cre-eGFP expression in LH astrocytes; quantification of orexin neurons (Mann-Whitney test: naïve= 16 (N=4), Cre-eGFP injected = 19 (N=5), *p*=0.381), quantification of s100β astrocytes (Mann-Whitney test naïve = 17 (N=6), Cre-eGFP injected = 18.40 (N=5), *p*=0.495), cellular identity of Cre-eGFP cells (Mann-Whitney test: s100β+ = 16.67 (N=6), Orexin-A+ = 0.60 (N=5), p <0.0001), and expression specificity of 94.5% in s100β+ astrocytes. **C**) Representative LH astrocytes expressing control virus (AAV2/5-GfaABC_1_D-eGFP) or AAV2/5-GfaABC_1_D-Cre-eGFP in 3D view and 2D slice view of branching structures (branch points & branching processes), 3D volume scale bar=5um, 2D slice scale bar=2um. **D**) Volume of -eGFP control or cre-expressing LH astrocytes from ELS male mice (unpaired *t*-test: eGFP = 4803 (N=3, n=19 cells), Cre = 4365 (N=3, n= 19 cells), *p*=0.491). **E**) Total branch points / volume (unpaired *t-*test: eGFP = 18.18 (N=3, n=19 cells), Cre = 31.91 (N=3, n =19 cells), *p*=0.007) **F**) Normalised Cx43 fluorescence intensity (a.u) in LH of - eGFP control or cre-astrocytes (unpaired *t*-test: eGFP = 71.37 (N=3, n= 14 cells), Cre = 102.6 (N=3, n=15 cells), *p* = 0.036). **G**) Volume of -eGFP control or cre-expressing LH astrocytes from ELS female mice (Mann-Whitney test: eGFP = 4622 (N=3, n=21 cells), Cre = 3545 (N=3, n=16 cells), *p*=0.130). **H**) Total branch points / volume (unpaired *t*-test: eGFP = 15.15 (N=3, n=21 cells), Cre = 31.06 (N=3, n= 16 cells), *p*=0.0018). **I**) Normalised Cx43 fluorescence intensity (a.u) in LH of -eGFP control or cre-astrocytes (unpaired *t*-test: eGFP = 91.32 (N=3, n=15 cells), Cre = 158.5 (N=3, n=15 cells), *p*<0.001). **J**) Experimental timeline for combined expression of AAV2/5-GfaABC_1_D-Cre-eGFP and AAV2/8-miniHCRT-tdTomato in *Nr3c1-*ELS mice, scale bars= 25um. **K**) Representative firing rates of orexin neurons in eGFP and Cre injected male mice and restoration of RMP and firing rates in Cre injected ELS mice (RMP: one-way ANOVA: *F*=10.32, *p*<0.001, Firing rate: one-way ANOVA: *F*=5.78, *p*=0.0072). **L**) Representative firing rates of orexin neurons in eGFP and Cre injected female mice and restoration of RMP and firing rates in Cre injected ELS mice (RMP: one-way ANOVA: *F*=7.90, *p*=0.0014. Firing rate: one-way ANOVA: *F*= 8.32, *p=*0.001). **M**) Running wheel behaviour of eGFP and Cre injected male mice: distance ran per day, total distance (one-way ANOVA: *F*=1.96, *p*=0.173), and light cycle activity (one-way ANOVA: *F*=3.23, *p*=0.001). **N**) Running wheel behaviour of eGFP and Cre injected female mice: distance ran per day, total distance (one-way ANOVA: *F*=6.02, *p*=0.011), and light cycle activity (one-way ANOVA: *F*= 1.24, *p*=0.315) N= mice. Ns, not significant, *P < 0.05, **P < 0.01, ***P < 0.001.

We first assessed the influence of astrocytic GR^KO^ on cell morphology in the LH using the same low-titre viral targeting approach as in **Fig. 2F**, but found no significant effect of astrocytic GR^KO^ on total volume of astrocytes (**Fig. 5D,G**). However, we saw a striking increase in the number of branch points / total volume, in both male and female ELS-GR^KO^ astrocytes (**Fig. 5E,H**). We next questioned whether this GR^KO^-induced morphological reorganisation influenced connexin-43 expression, one hallmark of this ELS model^40^. We observed a significant increase (43.8% in males, 73.5% in females) in normalised connexin-43 fluorescence in ELS-GR^KO^ astrocytes compared to control ELS-eGFP astrocytes (**Fig. 5F,I**). To ask whether this cellular rescue also attenuated the synaptic deficits (**Fig. 4**) we conducted stereotaxic surgery in ELS-*Nr3c1*^fl\fl^ mice for dual expression of AAV2/5-GfaABC_1_D-Cre-eGFP in LH astrocytes (green) combined with AAV2/8-miniHCRT-tdTomato to identify orexin neurons (red) (**Fig. 5J**). Not only did astrocytic GR^KO^ in the LH correct the ELS-induced changes in firing rate and RMP of orexin neurons, but it did so in a sex specific manner, attenuating the elevated firing rates in male ELS mice (**Fig. 5K**) and restoring reduced firing rates female ELS mice (**Fig. 5L**).

In light of these results, we conducted stereotaxic surgeries to induce astrocytic GR^KO^ in a separate cohort of ELS mice and subjected them to the two-week running wheel protocol to determine whether the synaptic rescue translated to behavioural recovery. Consistent with non-injected naïve and ELS litters (**Fig. 1M&N**), we observed consistent sexually dimorphic effects of ELS on running wheel behaviour in our control virus-injected (AAV2/5-GfaABC_1_D-eGFP) mice with light cycle hyperactivity in male mice (**Fig. 5M**) and hypoactivity in female mice (**Fig. 5N**). In agreement with the synaptic data, we report that astrocytic GR^KO^ in the LH rescued the effects of ELS on running wheel behaviour, diminishing light cycle hyperactivity in male ELS mice (**Fig. 5M**) and restoring distance ran per day in female ELS mice (**Fig. 5N**). Taken together, these data support the hypothesis that astrocytic GR signalling mediates the effects ELS on astrocyte morphology in the LH which in turn modulates intrinsic excitability of orexin neurons to perturb diurnal activity rhythms in a sex-specific manner.

## Discussion

In this study, we investigated the contribution of astrocytes in stress-induced behavioural change. We show that astrocytic stress signalling drives the cellular, synaptic, and behavioural effects of early-life stress on lateral hypothalamus-dependent behaviour in a sex-specific manner. Using an ELS paradigm shown to prompt long-term cellular and behavioural deficits, we report a strong sexual dimorphism in stress-induced activity changes with hypoactivity in female and hyperactivity in male mice accompanied by increased blood glucocorticoid levels in both sexes. Focusing on the lateral hypothalamus, we observed reorganisation of astrocytic morphology, downregulation of key astrocytic proteins, and disruption of intrinsic firing of orexin neurons. Importantly, we show that manipulating astrocyte function alone is *sufficient* to perturb diurnal patterns of wheel running using a chemogenetic strategy. Considering increased glucocorticoid levels was a conserved phenotype between sexes, we tested whether targeting this pathway could alleviate the effects of stress on cellular, synaptic, and behavioural scales. Using transgenic mice with astrocyte-specific downregulation of glucocorticoid receptors in the lateral hypothalamus, we reveal that stress signaling in astrocytes is *necessary* for the effects of ELS on cellular pathology, aberrant firing patterns of orexin neurons, and wheel running behavior in both female and male mice.

The precise ELS model employed in this study has been shown to have wide influence on transcriptomic profiles in both cortical and sub-cortical brain regions, with distinct sexual dimorphism in the transcriptional response to ELS^66^. In agreement, we report sex differences at both behavioural and synaptic levels. We focused our work on lateral hypothalamic astrocytes and their control of LH-specific behaviours however, ELS exerts its strong influence on behaviour via pleiotropic effects across multiple brain regions including the amygdala, PFC, hippocampus^40,101^. As early-life stress (or adversity) is one of the strongest risk factors associated with the development of anxiety and depression^102^ understanding the cellular mechanisms and what makes stress during development so profound is essential.

As almost all brain cell types express the *nr3c1* gene that codes for the glucocorticoid receptor^103^, the elevated blood corticosterone levels at P45 that we show and any subsequent signaling in the CNS is bound to have widespread effects. It would be of great interest to understand the specific contribution of GR signalling in other glial cell types^104,105^ and in neurons to determine their respective contributions to ELS-induced behavioural change. One reason why we believe astrocytes are particularly sensitive to glucocorticoids is due to the proximity of astrocytic endfeet to brain cerebrovasculature^106^, the entry point of blood glucocorticoids to the brain parenchyma enabling them to coordinate synaptic signals with signals from the vasculature^107^ including changes in circulating glucocorticoids. Indeed, RNA-seq data suggests a 7-fold higher expression of *nr3c1* in astrocyte (28.5 FPKM) compared to neurons (3.75 FPKM)^103^, so we postulate astrocytes are spatially and transcriptionally poised as primary mediators of the central effects of glucocorticoids. This notion is supported by the data presented here which reveals astrocytic GR signalling as a key mediator of the effects of early-life stressors. This emphasises the need for further understanding of astrocytic GR signalling in both physiological and pathological conditions.

We observed distinct sexual dimorphism in the behavioural and synaptic response to ELS. A noteworthy finding from our patch-clamp experiments is that both male and female ELS orexin neurons display a persistent depolarisation of the RMP yet, ELS has opposing effects of firing rates, driving behaviour. This discordance points to distinct mechanisms driving the synaptic phenotypes in ELS. Given that local (LH specific) deletion of astrocytic GR signalling corrected the sex-specific effects of ELS on both orexin neuron firing rates and behaviour, we propose the mechanism underlying ELS originates from a common dysfunction of astrocytes with sex differences downstream of initial glucocorticoid signalling.

Our ELS paradigm coincides with peak astrocyte process outgrowth and integration into synapses (P14-21), which could explain the atrophic phenotypes observed in both ELS male and female astrocytes in the LH^108^. Furthermore, there is evidence for differences in gene expression profiles across post-natal development between male and female astrocytes^12^. This could explain the more pronounced effect of ELS on female astrocyte morphology compared to males, where female astrocytes exhibit an arrested development as a consequence. Therefore, it is plausible this ELS paradigm functions as a “two-hit susceptibility hypothesis”, where ELS alters post-natal development of astrocyte morphology (Hit 1 – *primer*). Stress induced elevations in glucocorticoids activate signaling pathways resulting in dysfunctional astrocyte-synapse interactions and behaviour (Hit 2 – *effector*). To truly test whether disrupted astrocyte morphology in ELS is one of the main drivers and/or primers of the synaptic and behavioural dysfunction, the field requires more precise genetic tools to manipulate astrocyte morphology in real-time and measure its effects on brain circuits underlying behaviour.

Globally, our work contributes to the growing body of literature demonstrating the important role of glial cells in behavioural regulation and how this can be perturbed in pathology. Showing that astrocyte dysfunction is both necessary and sufficient to drive stress-induced behavioural change, supports a shift in how to conceptualise the cellular and molecular mechanisms of stress. The responsiveness of astrocytes to hormones is not limited to glucocorticoids but has also been shown for insulin ^109–111^, leptin ^112^, ghrelin ^113^. Disruption of these hormone signalling pathways in astrocytes also affected behaviour including wheel running^114^. Together these independent observations underscore the important role of astrocytes in contextual guidance of neural circuits^115^, whereby astrocytic integration of both hormonal and synaptic information is key to adaptation. Our findings pave the way to a more comprehensive incorporation of astrocyte biology in behavioural neuroscience, towards multicellular models of brain function.

## Acknowledgements

We thank Aurélie Cleret-Buhot (CRCHUM cellular imaging core) for microscopy training, and the staff of the animal facility at the CRCHUM. Fonds de Recherche du Québec – Santé (FRQS; 296562 & 309889), CHUM Foundation, Fondation Courtois grants to C.M-R., and the Réseau Québécois sur le Suicide les troubles de l’humeur et les troubles Associées (RQSHA) pilot grant to L.R.D-H and C.M-R. L.R.D-H. was supported by a FRQS doctoral award. B.R. was supported by a Canada Graduate Scholarship Master’s award, and a recruitment award from Neuroscience Dept. Université de Montréal. C.M-R. was supported by a Junior 1 Chercheur-Boursier salary award from FRQS.

## Author Contributions

L.R.D-H, and C.M-R designed the study. L.R.D-H. carried out behavioural experiments, immunostaining and analysis of subsequent data. L.R.D-H. and S.H. carried out morphological analysis. L.R.D-H took blood samples and performed ELISA experiments. L.R.D-H. and A.B. carried out injection of viral vectors. L.R.D-H. and B.R. carried out electrophysiological experiments and L.R.D-H carried out analysis of subsequent data. S.P. maintained and generated mouse colonies. S.F. providing running wheel equipment and expertise. L.R.D-H and C.M-R wrote the manuscript. All authors contributed to ELS paradigm and approved the final version of the manuscript.

## Methods

### Animals

All experiments were performed in accordance with the guidelines for maintenance and care of animals of the Canadian Council on Animal Care (CCAC) and approved by the Institutional Committee for the Protection of Animals (CIPA) at the Centre Hospitalier de l’Université de Montréal. Both male and female C57BL/6J mice (Jax #000664) and Nr3c1^flx/flx^ (Jax # 021021) were used in the study, housed on a 12hr:12hr light:dark cycle (lights on at 06:30) with *ad libitum* access to food and water.

### Early Life Stress protocol

C57BL/6J or Nr3c1^flx/flx^ pups were separated from their mothers for 4-hours per day (ZT2-ZT6) between post-natal days10 and P17. During separation pups and mothers were placed into new cages with 70% less bedding. Bedding was weighed on P10 and divided into each cage (home cage, pup separation cage, and mother separation cage). Separated pups and mothers had *ad libitum* access to food and water and pup cages were placed on a water heating pad maintained at 34 degrees Celsius. After the final day of separation (P17) bedding was returned to the original amount and pups were housed with their mothers until weaning at P21.

### Behavioral assays

#### 2 week running wheel protocol

Mice were single housed with a low-profile wireless running wheels (Med Associates, NV-047 & -047V) for 2 weeks (1 week habituation, 1 week data acquisition). Mice were undisturbed for the 2 weeks with *ad libitum* access to food and water. Total distance was calculated with an inhouse MatLab script using ZT binned revolutions per day and running wheel circumference (2 x ν x r). Light cycle activity counts were averaged across 3 days during data acquisition (days 9-11). Curve analysis for 24-hour activity cycles were calculated from extracted 5-minute bin data and the following measures were quantified; Area Under the Curve (AUC), Peak wheel running, and Full-Width at Half-Max (FWHM).

#### Sucrose Preference Test (SPT)

Mice were housed with dual-water bottle AllenTown cages, for 48 hours with two bottles containing water (switched at 24 hours) to account for novelty induced water intake. Bottles were next replaced with either water or a 1% sucrose solution for 2 days, which were switched at 24 hours after sucrose presentation. Sucrose preference ratios were calculated using the following equation.

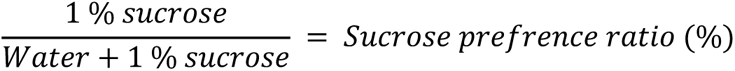

### Blood collection & Corticosterone analysis

Trunk blood was collected via rapid decapitation at ZT2 (08:30) or ZT14 (20:30). To minimise handling induced elevations in corticosterone mice were housed with clear plastic tubes used to move individual mice into an enclosed chamber filled with 5% isoflurane for 2 minutes. Mice were placed in enclosed Isoflurane chambers until loss of toe pinch reflect (maximum elapsed time of 2 minutes). Mice were decapitated and trunk blood was collected into BD 365963 Microtainer® Capillary Blood Collection Tubes and placed onto Ice. Blood was centrifuged for 5 minutes at 4 degrees Celsius at 5000rpm. Serum was aliquoted and stored at -80 degrees Celsius. Corticosterone measurements were obtained using an ENZO ELISA kit (ADI-900-097).

### Acute brain slice preparation

Coronal slices containing lateral hypothalamus were obtained from mice between P45-70. Animals were anesthetized with isoflurane and the brain was rapidly excised and placed in ice-cold NMDG based cutting solution saturated with 95% O_2_ and 5% CO_2_ containing the following (in mM): 119.9 NMDG, 2.5 KCl, 25, 1 CaCl2, 1.4 NaH2P04 and 20 D-glucose saturated with 95% O2 and 5% CO2. Slices (250µm thick) were cut on a vibratome (Leica VT1200, Nussloch, Germany). Slices were transferred to oxygenated NMDG cutting solution artificial CSF (aCSF) at 33 ± 0.5°C for 15 min followed by a 1-hour recovery period in artificial CSF (aCSF) at RT containing the following (in mM): 130 NaCl, 2.8 KCl, 1.25 NaH2PO4, 1.2 MgCl2, 2.5 CaCl2, 300-305 mOsmol. For experiments, slices were transferred to a recording chamber where they were perfused (2 ml/min) with aCSF at RT for the course of the experiment. Slices were used for a maximum of 5 hr after cutting.

### Electrophysiology

Orexin neurons in the LHa were identified through positive expression of AAV2/8-miniHCRT-TdTomato with a plan n 4x/0.10 magnification (Olympus) or LUMPlanFL N 40x/0.80 magnification (Olympus) objective with an upright scientific scicam pro camera. Whole-cell patch-clamp recordings of orexin neurons were performed in current- and voltage-clamp configuration. Recordings were made at room temperature. Data were acquired using a Multiclamp 700B amplifier (Molecular Devices) and digitized using a Digidata 1440A digitizer and pClamp/Clampfit 10.7 (Molecular Devices). Pipettes (from borosilicate capillaries; World Precision Instruments) had resistances of 4-8 mega-ohms and were filled with an internal solution containing the following (in mM): 105 K-gluconate, 30 KCl, 10 phosphocreatine, 4 ATP-Mg, 0.3 GTP-Tris, 0.3 EGTA (adjusted to pH7.2 with KOH ; 290-300 mOsmol) . For AAV2/8-miniHCRT-TdTomato validation experiments biocytin (2 mg/mL; Sigma) was added to the internal solution (**Supplemental Fig.3A&B**). All recordings were analyzed with Clampfit 10.2 and 10.6 from pClamp software (Molecular Devices). Liquid Junction Potentials were not adjusted for. Recordings were discarded if access resistance (Ra) values changed by more than 30% across recording time and/or whole-cell patch clamp recordings had Ra values higher than 25 mega-ohms. Action potential firing rate was calculated by automated analysis through Clampfit 10.2. Excitatory Post-Synaptic Potentials (ePSCs) were obtained in the voltage-clamp configuration (-70mV) for 2 minutes. For all ePSC analyses experimenters were blinded when scoring traces by hand. Input resistance (Rin) was determined by measuring the slope of the linear portion of the current-voltage (I-V) curve.

### Stereotaxic surgery

P30 mice were given a subcutaneous (*s.c.*) injection of Carprofen (20 mg/kg) minumum 1h prior to surgery. Animals were deeply anesthetised (5% isoflurane) before placing them into a stereotaxic frame (maintained with 1-2% isoflourane) (*Kopf instruments*). Surgical site was infiltrated with bupivacaine/lidocaine (2mg/kg *sc.*) 10 min prior surgery. Two small holes were made to bilaterally expose the brain surface under the skull. A 10ul Neuro syringe (Hamilton) was filled with the following AAV constructs produced at the Canadian Neurophotonics Viral Vector Core: AAV2/5-gfaABC_1_D-eGFP (1.5×10^11–12^ GC/ml), AAV2/5-gfaABC_1_D-Cre-eGFP (1.5×10^12^ GC/ml), AAV2/5-MiniHCRT-TdTomato (7.0×10^12^ GC/ml), AAV2/5-gfaABC_1_D-HM3d(Gq)-mCherry (1.5×10^12^ GC/ml), AAV2/5-gfaABC_1_D-HM4d(Gi)-mCherry (1.5×10^12^ GC/ml), AAV2/5-gfaABC_1_D-mCherry (1.5×10^12^ GC/ml). 600nl of vector per hemisphere was injected at 200nl/min into the lateral hypothalamic area (AP/DV/ML = ±1.6/-5.3/ ±0.9 mm) using a microinjection syringe pump (UMP3T-2 World Precision Instruments). For morphological analysis of astrocytes in the LHa viral titres were dropped ten-fold to 10^11^ GC/ml. Mice were given minimum 2 weeks to recover from surgeries before experiments.

### Chemogenetics

Male and female mice injected with either AAV2/5-gfaABC_1_D-HM3d(Gq)-mCherry, AAV2/5-gfaABC_1_D-HM4d(Gi)-mCherry, or AAV2/5-gfaABC_1_D-mCherry were singled housed for two weeks with running wheels (Methods: *2 week running wheel protocol*). Water intake was measured for the three last days of week 1 (days 4-7). Clozapine-N-Oxide (CNO) (Tocris, #4936) was standardised to body weight on day 7 and water intake averaged across days 4-7 and delivered into the drinking water for 3 days. CNO was delivered to mice at 1.25mg/Kg, 2.5mg/Kg, and 5mg/Kg.

### Brain processing

Mice were anaesthetised in an enclosed chamber filled with isoflurane (5% for induction, 3-4% for maintenance, v/v) until loss of toe pinch reflex. A transcardial perfusion was used to ensure uniform preservation of tissue and brains were fixed in 4 % paraformaldehyde (PFA) for 24 hours at 4 degrees Celsius. The fixed brains were then placed in 15ml falcon tubes containing 30% sucrose solution for a minimum of two days. Brains were embedded in optimal cutting temperature (OCT) compound and flash frozen in 2-methylbutane between -40 and -50 degrees Celsius and then stored at -80 until cryosectioning. Brains were cut at 35µm thickness at -20 degrees Celsius using a Leica CM3050S Cryostat. Slices were stored in a cryoprotector solution at 4 degrees Celsius until immunohistochemistry.

### Immunohistochemistry

Free floating brain slices were washed 3 times in 1X PBS for 15 minutes to remove cryoprotector solution. Slices were permeabilised in block-perm solution (3% Bovine-Serum-Albumin, 0.5% Triton^10%^ in PBS) for a duration of 1 hour. Slices were incubated with antibodies for 24 hours at 4 degrees Celsius. Slices were incubated with primary antibodies; [Rabbit] anti-s100β (1:1000, Abcam, ab52642), [Mouse] anti-glucocorticoid receptor (1:500, ThermoFisher, MA1-510), [Chicken] anti-GFAP (1:1000, ThermoFisher, PA1-10004), [mouse] anti-Cx43 (1:1000 ThermoFisher, 3D8A5), [mouse] anti-orexin-A (1:1000, NovusBio, MAB763) and [rabbit] anti-eGFP (1:500, ThermoFisher, G1362). After primary antibody incubation slices were washed 3 times in 1X PBS for 15 minutes to remove non-specific binding. Slices were incubated with secondary antibodies in a DAPI solution (1:10000) at RT in aluminium foil for 1 hour. Slices were incubated with secondary antibodies; [goat] anti-rabbit Alexa 488 (1:1000, Jackson Immuno Research, 111-545-144), [goat] anti-mouse Alexa 647 (1:1000, ThermoFisher, A32728), [goat] anti-chicken Alexa 568 (1:1000, ThermoFisher, A11041). After secondary antibody incubation slices were washed again 3 times in 1X PBS for 15 minutes before being mounted onto Fisherbrand microscope slides using ProLong™ Glass Antifade mountant (P36982).

### Microscopy

Slices were imaged using a Leica TCS SP5 laser scanning confocal microscope with oil immersion Plan-Apochromat 63x objective 1.4 NA,. 16-bit images of 246 x 246 um areas were acquired at 400Hz (frame size (x*y); 1024 x 1024, pixel size; 250nm). 25-30 µm z-stacks were acquired with a step size of 0.5 µm. 15µm max intensity z-projections of the lateral hypothalamus were analysed using Image-J (Fiji) to obtain A.U fluorescence intensity measures of secondary antibodies attached to primaries with specificity to epitopes; GFAP, Cx43, and GR. We centered astrocytes with a 2241.025µm^2^ ROI and applied a fluorescence threshold for GFAP and Cx43 fluorescence and measured integrated density A.U (thresholds; default & Otsu respectively). AU measurements were standardised to ROI sizes to obtain normalised fluorescence values. For Astrocytic cell counts s100β staining masked over DAPI channels and we used the analyse particles function on Fiji, counting the number of DAPI nuclei larger than 10um in diameter. Orexin cells were counted manually using the Fiji pointer tool. For nuclear Glucocorticoid Receptor measures, astrocyte (s100β+) DAPI nuclei were thresholded and used as ROIs for nuclear measures of GR fluorescence (s100β^+^ + DAPI^+^).

#### Astrocyte Morphology

Mice were bilaterally injected with a low titre AAV2/5-gfaABC_1_D-eGFP into the LHa (Methods: stereotaxic surgery). After 3 weeks mice were transcardially perfused and fixed for immunohistochemistry. To obtain singular astrocytes for morphological reconstruction we imaged astrocytes at injection site periphery regions to limit overlap of eGFP+ expressing astrocytes whilst remaining in LHa brain territory. Astrocytes were imaged using a LSM laser scanning confocal microscope with oil immersion Plan-Apochromat 63x objective 1.4 NA,. 16-bit images of 101.4 x 101.4 µm areas were acquired at 6 fps (frame size (x*y); 1724 x 1724, pixel size; 0.06µm). 25-30µm z-stacks were acquired with a step size of 0.5µm. Isolated astrocytes were reconstructed using the IMARIS platform where we calculated branching, process lengths, 3D volumes, and sholl analyses.

### Statistical analyses

Results are presented as mean ±S.E.M. or box and whisker plots for data with one variable and were analyzed with the two-tailed Student’s *t* test or Mann-Whitney test respectively. Chemogenetic experiments used repeated measures *t-*tests to non-parametric equivalents. All data was tested for normality and data sets with more than two conditions were first screened for a Gaussian distribution with Kolmogorov-Smirnov test followed by analysis either with one-way/repeated measures ANOVA or Kruskal-Wallis/Friedman test when needed and Tukey’s multiple-comparison parametric *post hoc* test (data with Gaussian distribution) or by a Dunn’s multiple-comparison non-parametric *post hoc* test (data with non-Gaussian distribution). Diurnal running wheel distances were analysed using a two-way repeated measures ANOVA. Statistical tests are represented with the following significance thresholds, *, p < 0.05; **, p < 0.01 and ***, p < 0.001. All data were analyzed using GraphPad Prism software (Version 9, GraphPad, USA).

---

**Supplemental Figure 1.**
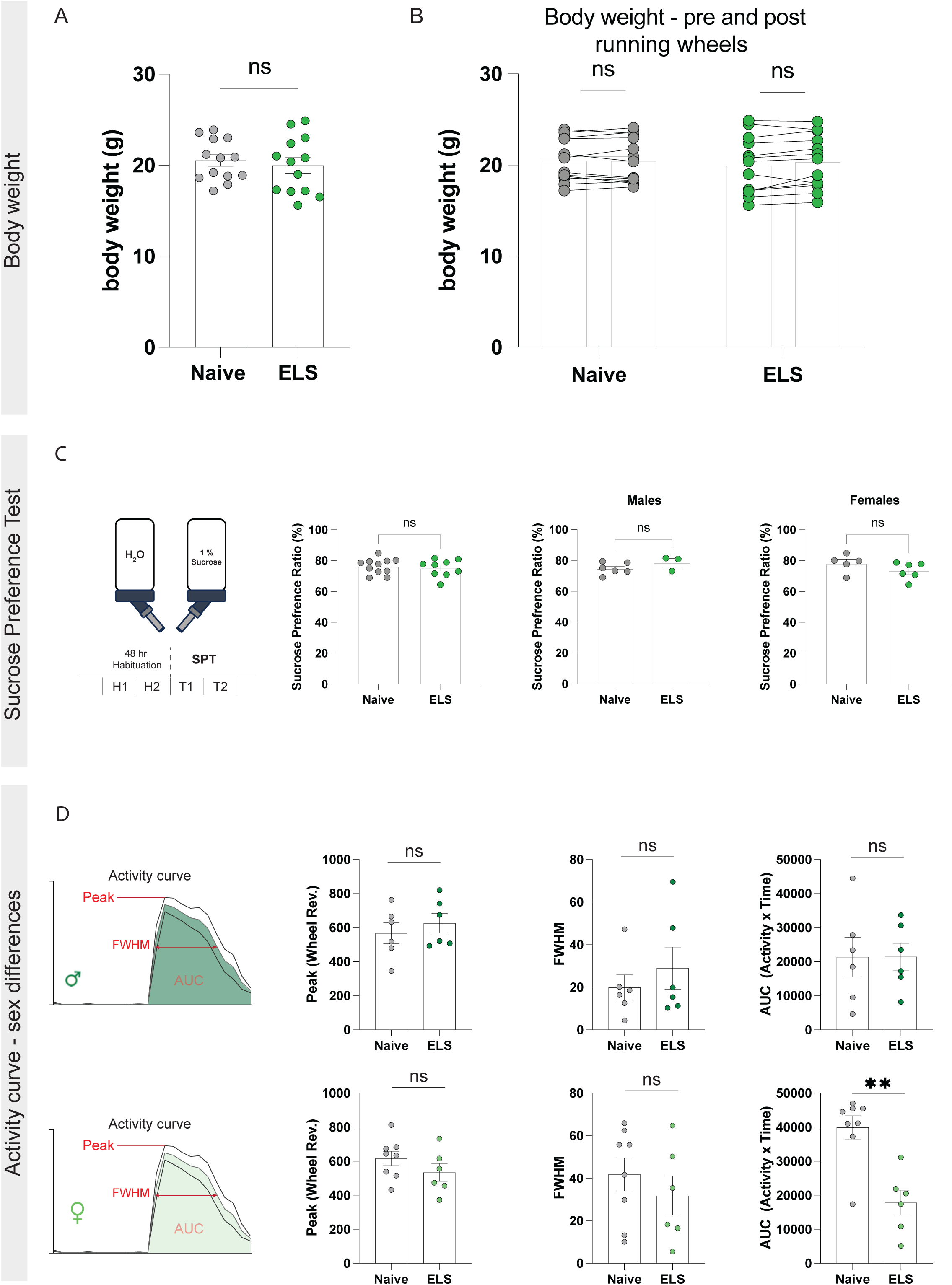
Behavioural phenotyping of naïve and ELS mice. **A**) Body weight (g) of naïve and ELS mice at beginning of experimental window, P45 (unpaired *t*-test: naïve= 20.53 (N=13), ELS=19.98 (N=13), *p*=0.610). **B**) Body weight of naïve and ELS mice pre- and post-2 weeks running wheel protocol (Naïve: paired *t*-test: *t*_12_=0.194, *p*=0.849. ELS: paired *t*-test: *t*_12_=1.903, *p*=0.081). **C**) Sucrose Preference Test in naïve and ELS mice (unpaired *t*-test: naïve=76.34 (N=11), ELS=75.19 (N=9), *p*=0.631) with no sex differences. Males: Mann-Whitney test: naïve=74.18 (N=6), ELS=80.98 (N=3), *p*=0.262, Females: unpaired *t*-test: naïve=78.28 (N=5), ELS=73.51 (N=6), *p=*0.199. **D**) Activity curve analyses for naïve and ELS mice split by sex. *Top*: male. Peak wheel running (Mann-Whitney test: naïve=576 (N=6), ELS=598 (N=6), *p=*0.485), FWHM (Mann-Whitney test: naïve=17.45 (N=6), ELS=17.63 (N=6), *p=*0.812), AUC (Mann-Whitney test: naïve=20927 (N=6), ELS=20373 (N=6), *p=*0.937). *Bottom*: female. Peak wheel running (Mann-Whitney test: naïve=639 (N=8), ELS= 515 (N=5), *p=*0.282), FWHM (Mann-Whitney test: naïve=48.74 (N=8), 26.89 (N=6), *p=*0.470). AUC (Mann-Whitney test: naïve=43853 (N=8), ELS=18974 (N=6), *p=*0.008). N= mice. Ns, not significant, *P < 0.05.

**Supplemental Figure 2.**
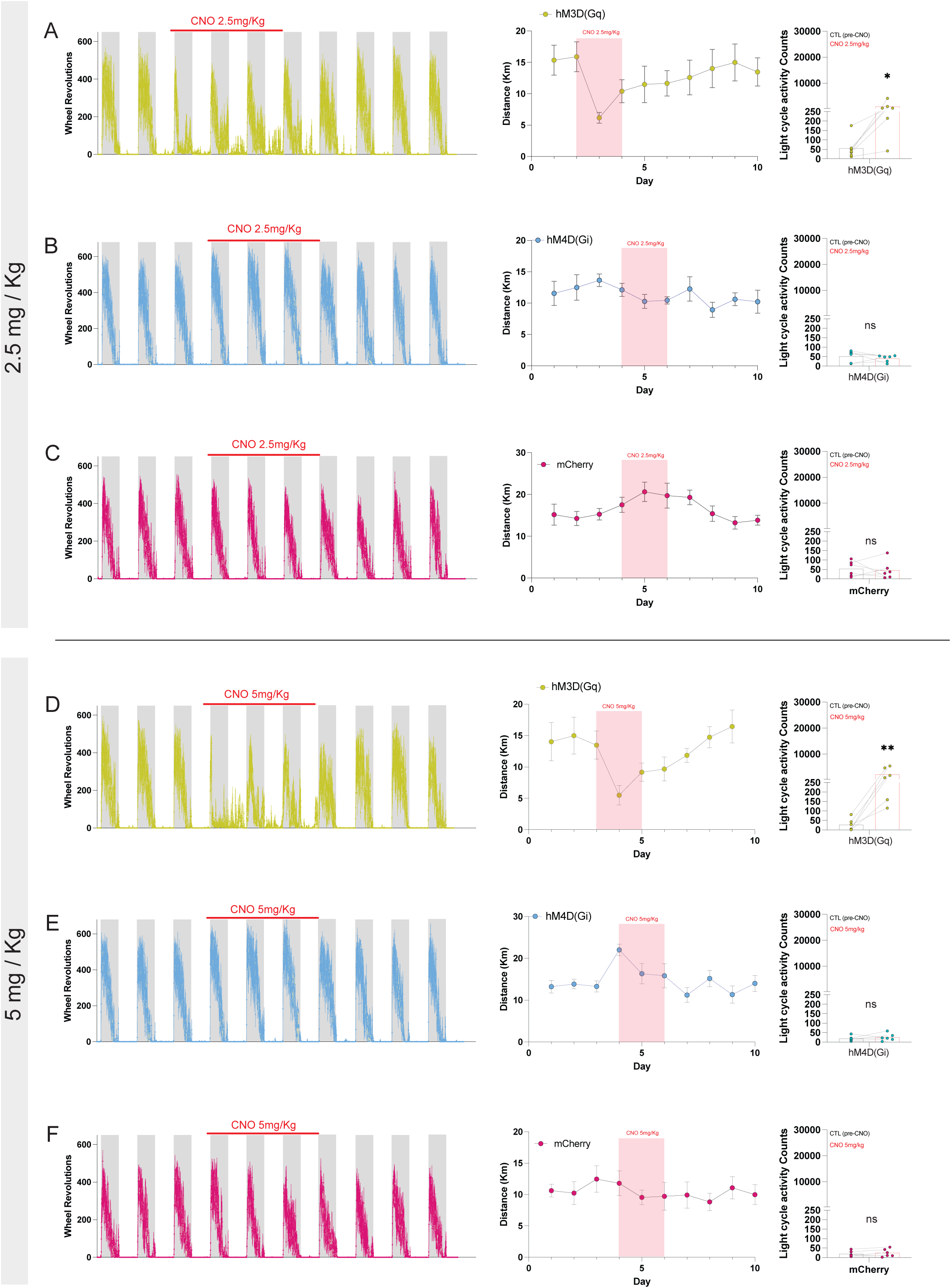
Astrocytic Gq-coupled calcium fluctuations in the LH perturb activity rhythms at ascending doses of CNO (2.5 and 5 mg/kg). **A-C)** Data for 2.5mg/kg CNO. **A**) Representative traces of running wheel activity from hM3D(Gq)-injected mice (N=6), distance ran per day, and quantification of light cycle activity pre and during delivery of CNO (ratio paired *t*-test: *t*5=3.81, *p*=0.0125). **B**) Representative traces of running wheel activity from hM4D(Gi)-injected mice (N=6), distance ran per day, and quantification of light cycle activity pre and during delivery of CNO (paired *t*-test: *t*5=0.803, *p*=0.458). **C**) Representative traces of running wheel activity from mCherry-injected mice (N=6), distance ran per day, and quantification of light cycle activity pre and during delivery of CNO (paired *t*-test: *t*5=0.312, *p*=0.768). **D-F)** Data for 5mg/kg CNO. **D**) Representative traces of running wheel activity from hM3D(Gq)-injected mice (N=6), distance ran per day, and quantification of light cycle activity pre and during delivery of CNO (ratio paired *t*-test: *t*5=5.915, *p*=0.002). **E**) Representative traces of running wheel activity from hM4D(Gi)-injected mice (N=6), distance ran per day, and quantification of light cycle activity pre and during delivery of CNO (paired *t*-test: *t*5=0.802, *p*=0.459). **F**) Representative traces of running wheel activity from mCherry-injected mice (N=6), distance ran per day, and quantification of light cycle activity pre and during delivery of CNO (paired *t*-test: *t*5=1.34, *p*=0.232). N= mice. Ns, not significant, *P < 0.05, **P < 0.01.

**Supplemental Figure 3.**
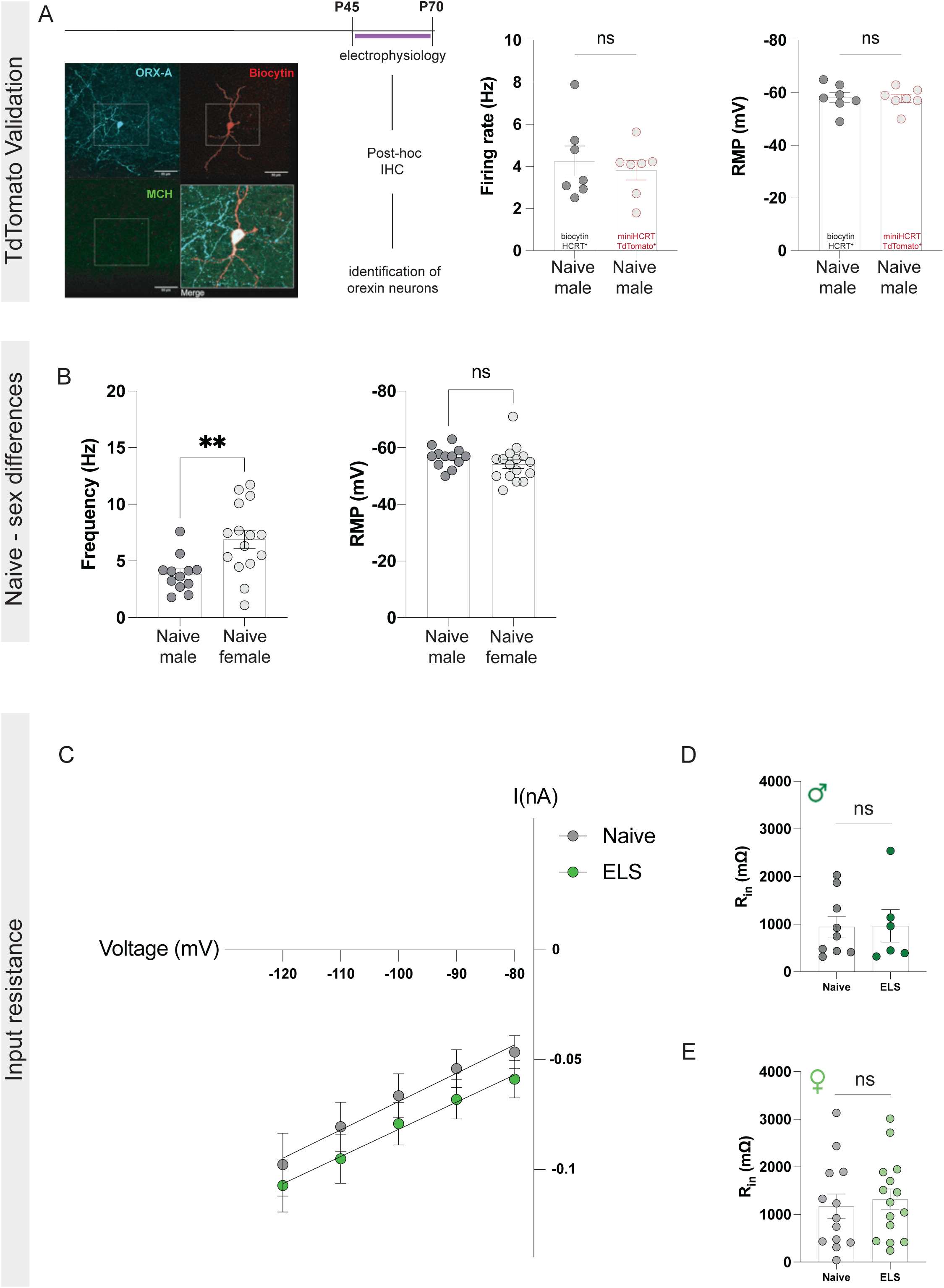
AAV2/8-mini-HCRT-tdTomato validation and intrinsic excitability. **A**) Comparison of tdTomato+ orexin neurons to biocytin filled orexin neurons identified through post-hoc IHC: firing rate (unpaired *t*-test: naïve male (biocytin)= 4.25 (N=3, n=7 cells), naïve male (tdTomato+)= 3.83 (N=3, n=7 cells), *p*=0.625) and resting membrane potential (unpaired *t*-test: naïve male (biocytin)= -58.19 (N=3, n=7 cells), naïve male (tdTomato+)= -57.83 (N=3, n=7 cells), *p*=0.889), scale bars=50um. **B**) Sex differences in basal firing rates of orexin neurons (unpaired *t*-test: naïve male= 3.85 (N=4, n= 12 cells), female naïve=6.91 (N=4, n= 15 cells), *p*=0.005) with no differences in resting membrane potential (unpaired *t*-test: naïve male=-56.65, naïve female =-54.19, *p*=0.226) **C**) Average current-voltage relationships in orexin neurons from naïve and ELS brain slices. **D**) Input resistance (R_in_) of orexin neurons from naïve and ELS male mice (unpaired *t*-test: naïve= 948.7 (N=3, n=9 cells), ELS= 965.7 (N=2, n=6 cells), *p*=0.965). **E**) Input resistance (R_in_) of orexin neurons from naïve and ELS female mice (unpaired *t*-test: naïve= 1173 (N=3, n= 13 cells), ELS= 1321 (N=3, n=15 cells), *p*=0.661). N= mice. Ns, not significant.

**Supplemental Figure 4.**
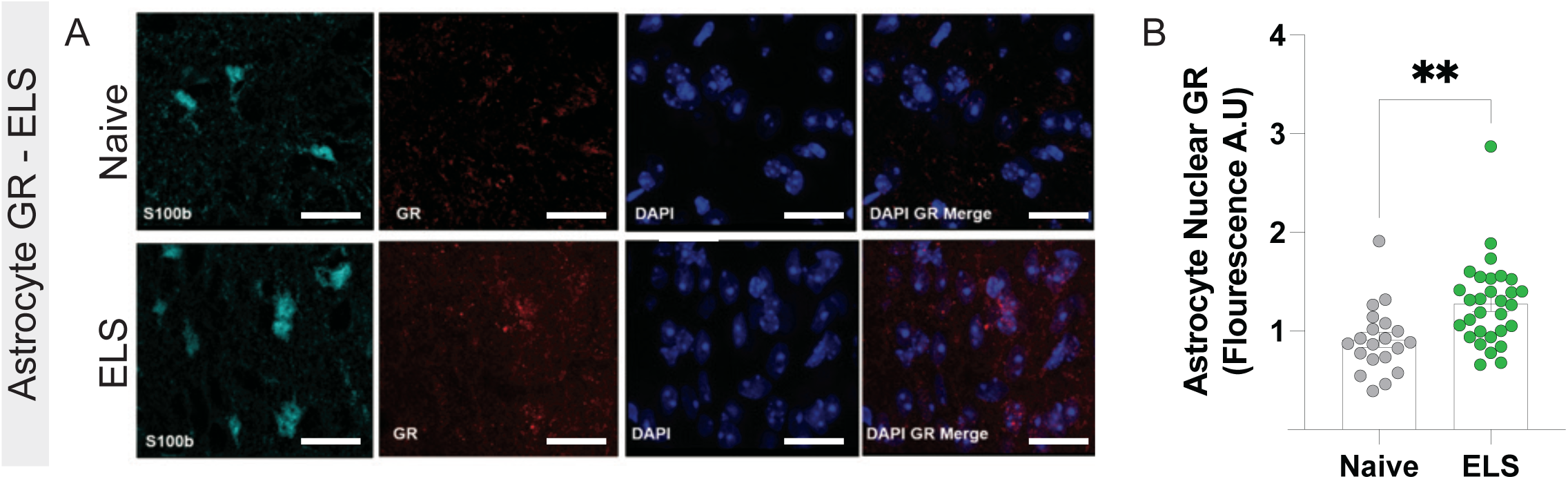
Elevated astrocytic glucocorticoid receptor signalling in ELS mice. **A**). Representative IHC of s100β+ astrocytes in the LH with co-staining against *Nr3c1 (*GR). **B**) Quantification of nuclear (DAPI-bound) GR fluorescence (a.u) in LH astrocytes (unpaired *t*-test: naïve= 9114 (N=4, n=20 cells), ELS= 12784 (N=6, n=30 cells), *p=*0.0026). N= mice. **P < 0.01.

**Supplemental Figure 5.**
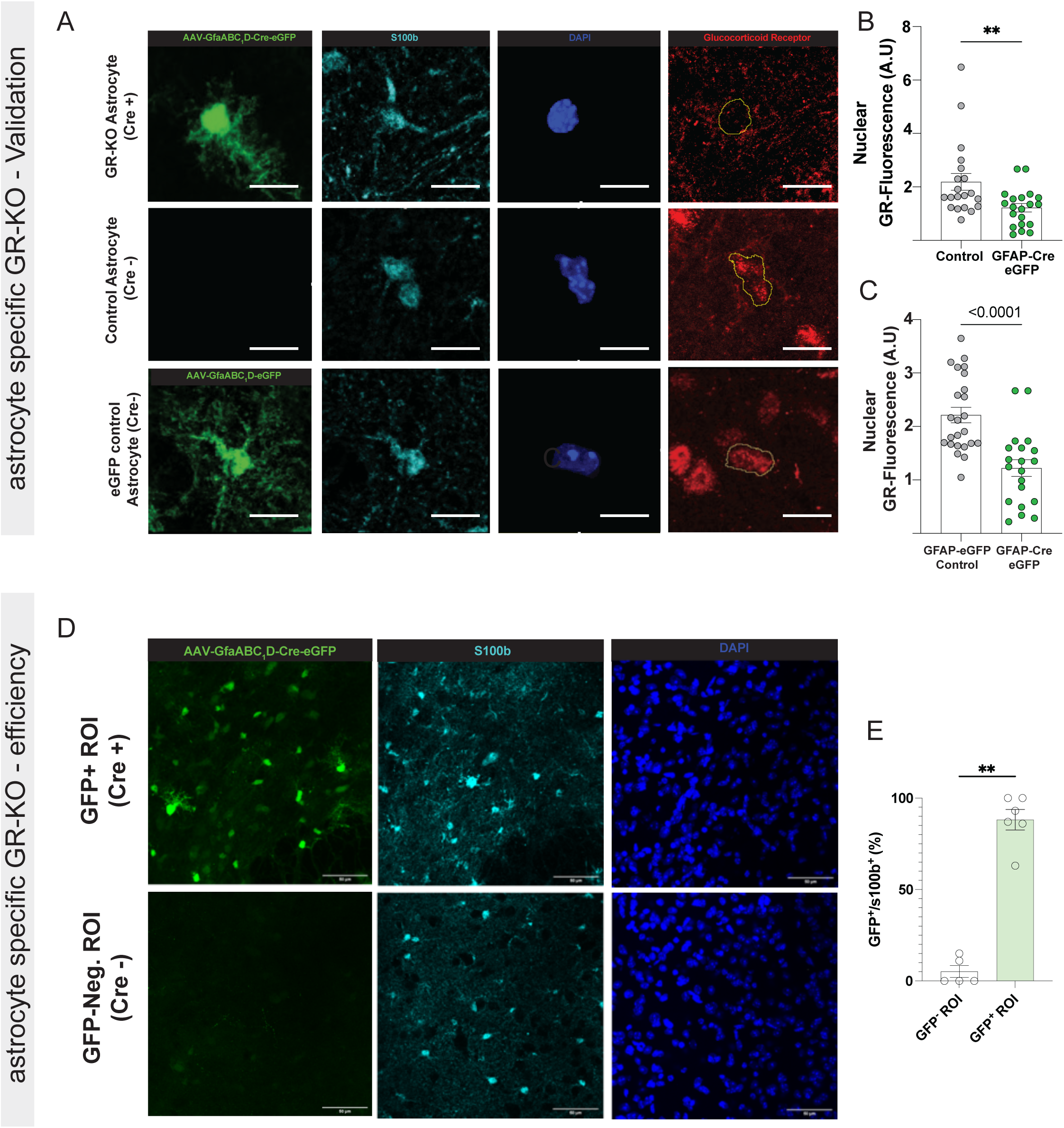
Viral validation of GR-KO in lateral hypothalamic astrocytes. **A**). Representative IHC of GR-KO astrocytes (AAV2/5-GfaABC_1_D-Cre-eGFP), within-brain control astrocytes (no virus), and eGFP control astrocytes (AAV2/5-GfaABC_1_D-eGFP) colocalized with astrocyte marker s100β and associated nuclear GR fluorescence, scale bar =25um. **B**) Quantification of nuclear GR fluorescence (a.u) in within-brain control and GFAP-Cre-eGFP (GR-KO) astrocytes (Mann-Whitney test: control=21898 (N=4, n=20 cells), GFAP-Cre-eGFP= 12207, (N=4, n=20 cells), *p=0.006*). **C)** Quantification of nuclear GR fluorescence (a.u) in GFAP-eGFP control and GFAP-Cre-eGFP (GR-KO) astrocytes (unpaired *t*-test: GFAP-eGFP= 22447 (N=4, n=20 cells), GFAP-Cre-eGFP= 12207, (N=4, n=20 cells), *p <0.0001).* **D-E**) AAV2/5-GfaABC_1_D-Cre-eGFP astrocyte specific expression efficiency (%) in GFP+ (Cre+) and GFP-(Cre-) ROIs, scale bars=50um (Mann-whitney test: GFP-ROI=0.05 (N=5), GFP+ ROI=90 (N=6), *p*=0.0043). N= mice. **P < 0.01.

## Statistics Table

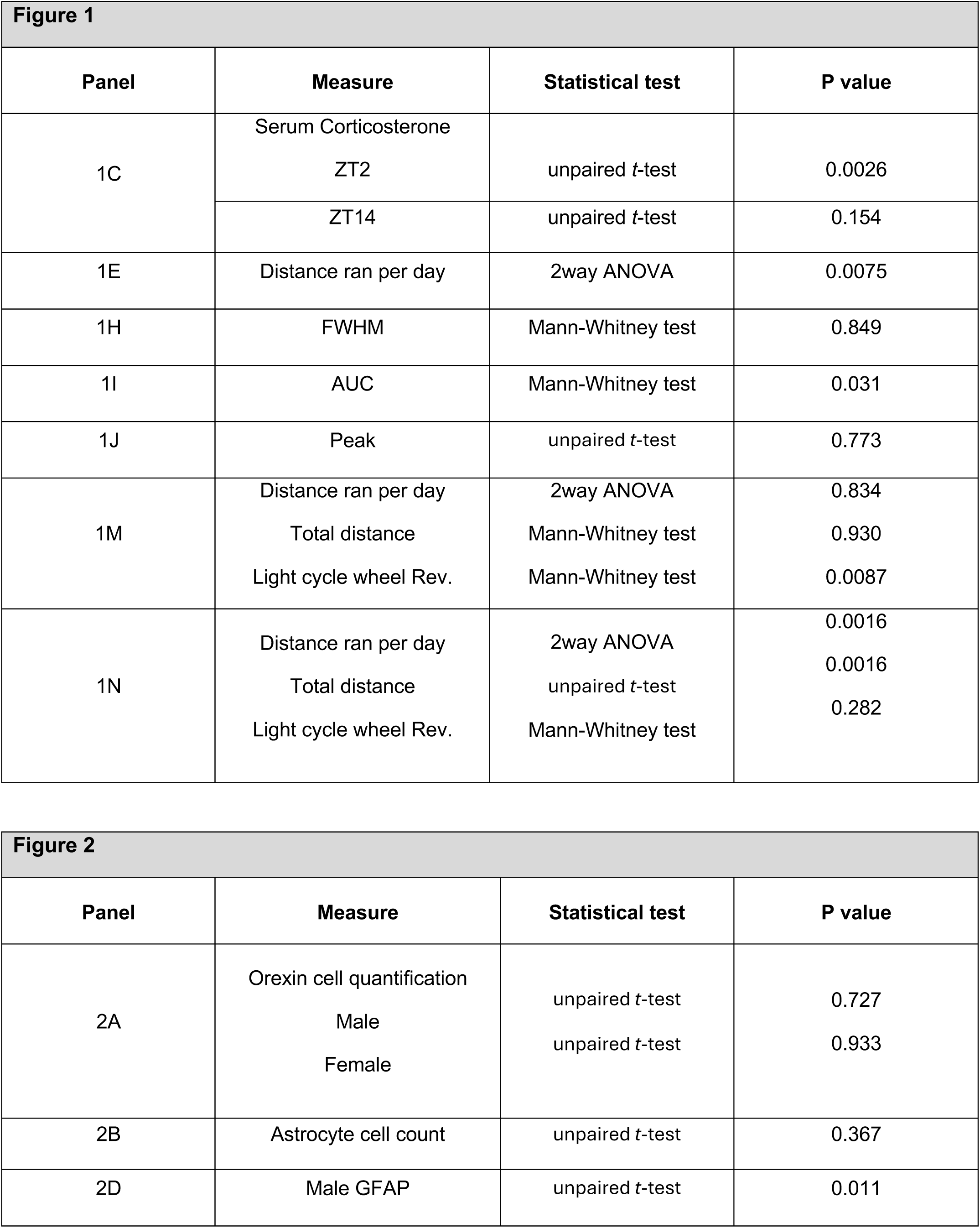
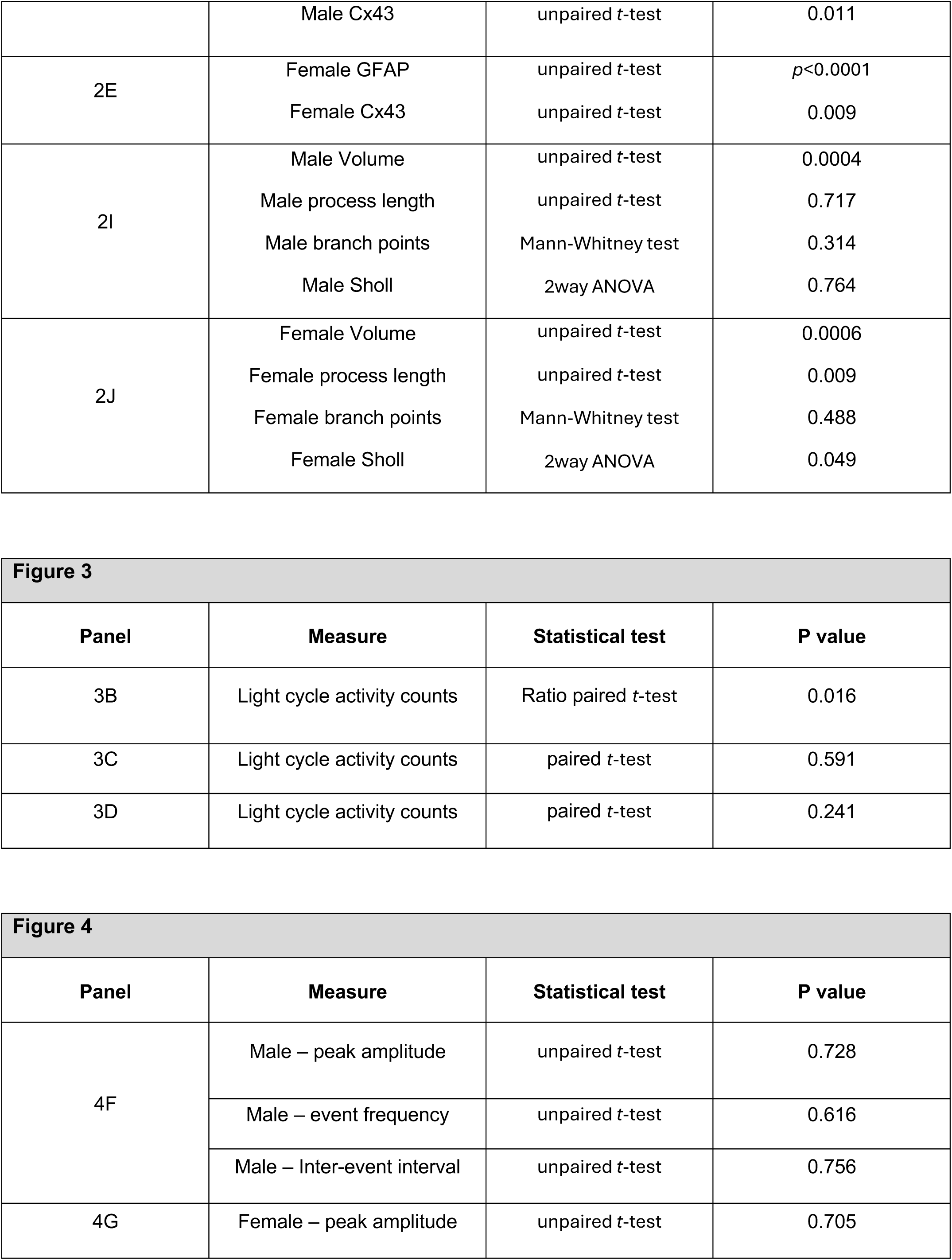
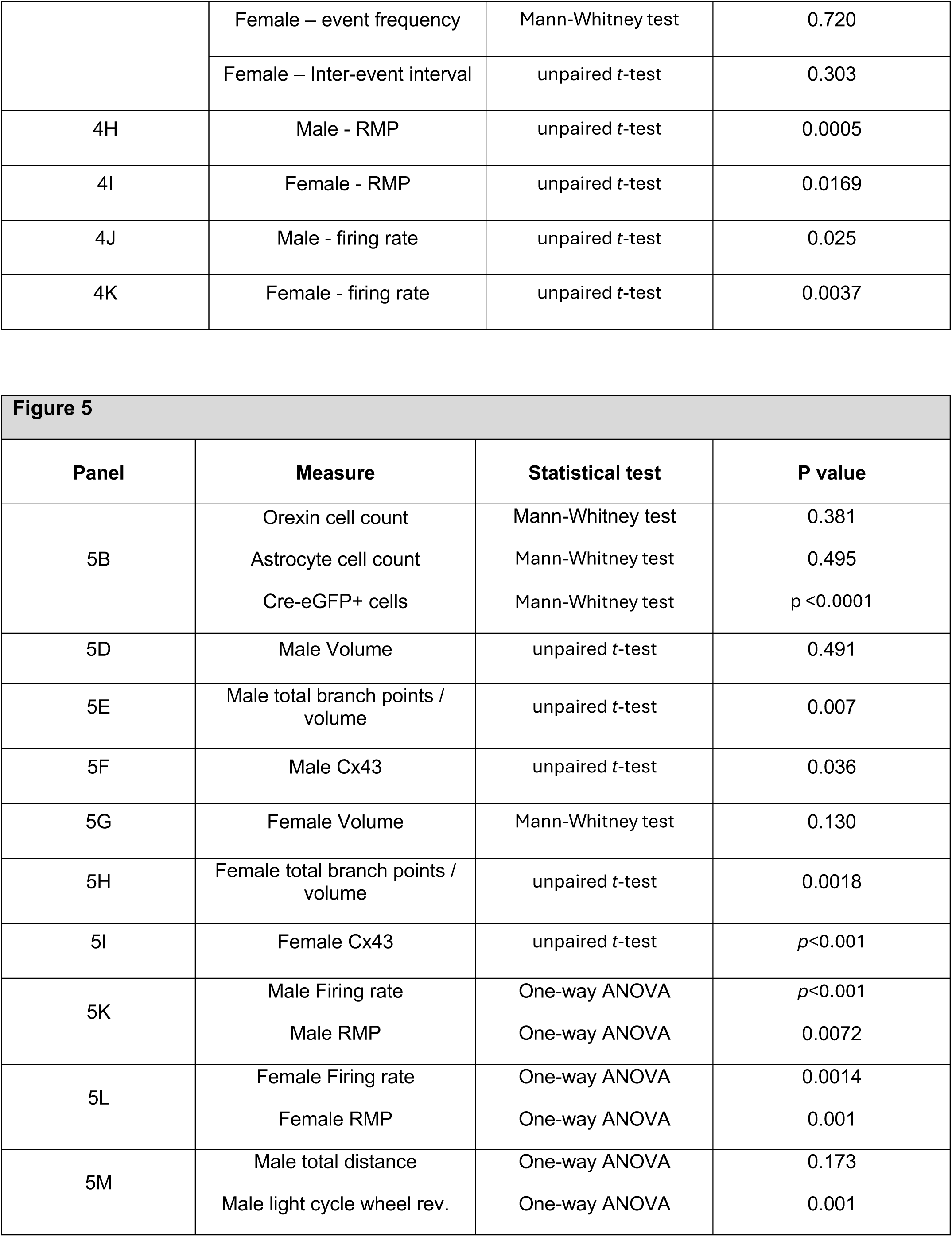
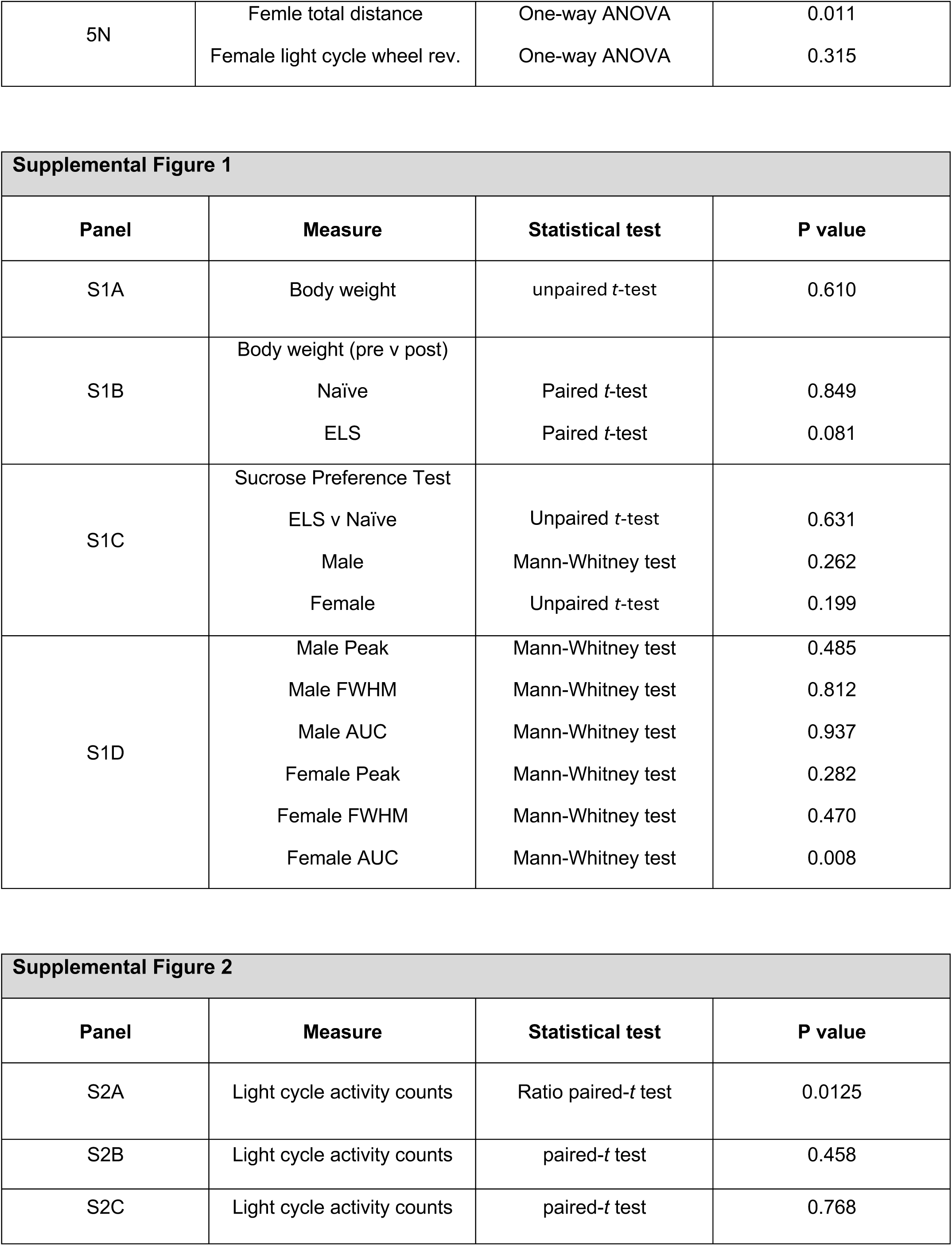
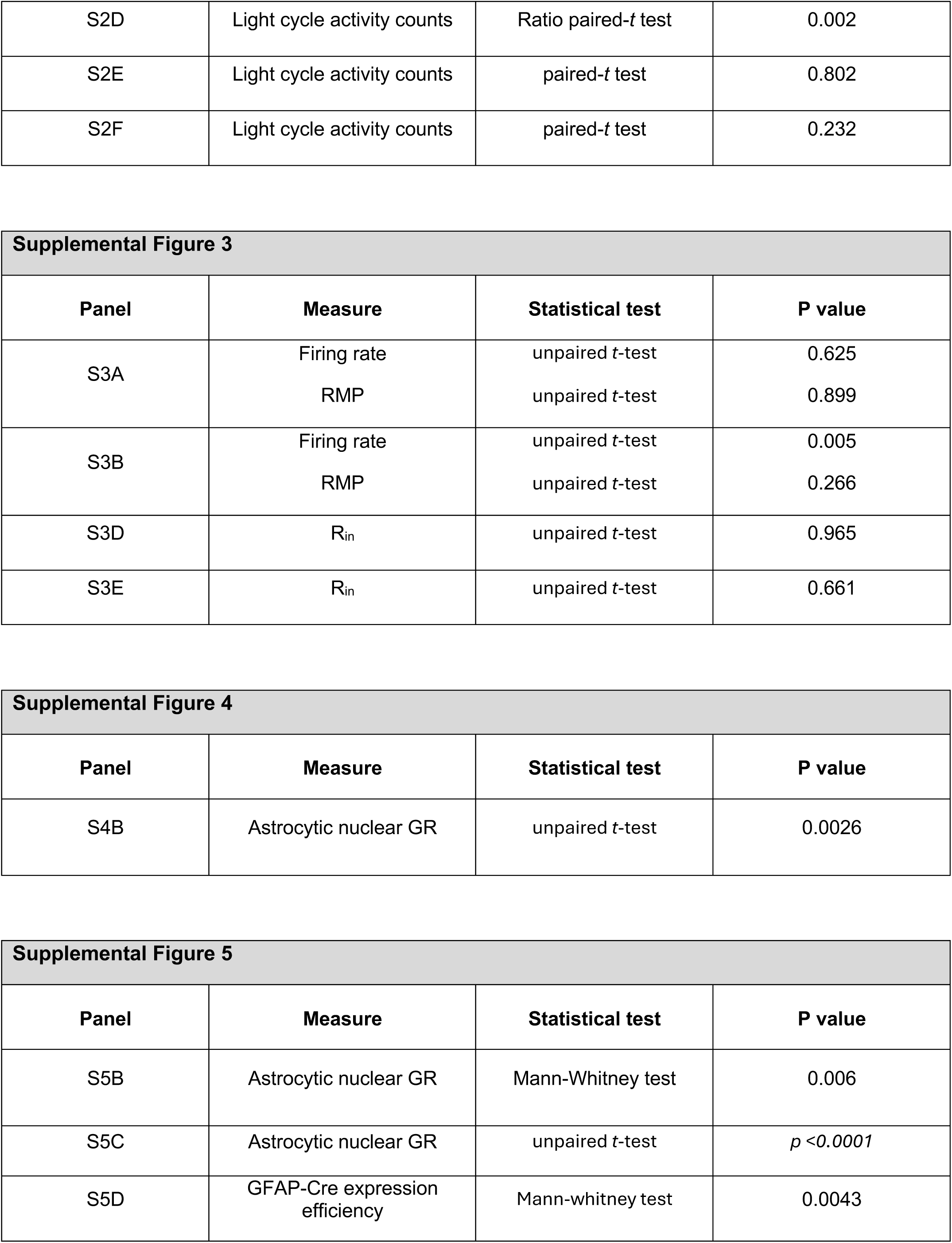

